# Enhancing adeno-associated virus cellular entry through receptor engineering

**DOI:** 10.64898/2026.05.26.727777

**Authors:** Bijay P Dhungel, Rajini Nagarajah, Cynthia Metierre, Yue Feng, Vivienne Kaiser, Harrison Bazley, Catherine Curry, Divya Gokal, Alex Sherman, Qian Peter Su, Mehdi Sharifi Tabar, John EJ Rasko, Charles G Bailey

## Abstract

Adeno-associated viruses (AAV) are approved for gene therapy of several genetic disorders; however, key aspects of AAV cellular entry remain poorly understood. We previously identified carboxypeptidase D (AAVR2) as an AAV receptor distinct from the multi-serotype AAV receptor KIAA0319L (AAVR). In this study, we investigate the molecular mechanisms and biological roles of AAVR and AAVR2 in mediating AAV gene transfer. Using proximity-dependent biotin identification (BioID), we defined the interactome of AAVR in the presence or absence of AAV8 and identified various proteins involved in viral entry including AAVR2. We confirmed a direct physical interaction between AAVR and AAVR2, mediated by non-AAV interacting regions in the C-termini. Further, we identified functional motifs within the carboxy-terminal tails of both receptors to facilitate the engineering of chimeric receptors with enhanced activity. Functional assays demonstrated that the overexpression of AAVR or AAVR2 enhances the cellular attachment and entry of AAV in a serotype-specific manner. Finally, we generated a stable cell line expressing a minimal functional AAVR2 with increased sensitivity for *in vitro* potency testing for AAVR2-engaging serotypes like AAV8. Collectively, these findings reveal significant functional similarities in AAV receptor biology and establish a framework for engineering receptor-guided modalities.

## Introduction

Adeno-associated virus (AAV) is a single-stranded, helper-dependent parvovirus widely used to deliver human gene therapies, having achieved multiple regulatory approvals and hundreds of advanced-stage clinical trials^1^. However, recent fatalities reported in multiple clinical trials following high dose AAV administration highlight critical gaps in the understanding of AAV transduction biology^1–5^. Although AAV capsids with altered cell/tissue tropism exist, molecular pathways facilitating cellular entry of AAV remain incompletely understood^1,6,7^.

AAV is internalized via receptor-mediated endocytosis followed by intracellular trafficking through the endosomes and Golgi to the nucleus^7^. Proteoglycans act as attachment factors increasing the surface-dwelling of AAV to facilitate interaction with proteinaceous receptors like KIAA0319L (or AAVR)^7,8^. AAVR is a highly conserved entry factor essential for the transduction of many, but not all, serotypes^9^. Known AAVR-independent transduction pathways and serotypes led to our discovery of carboxypeptidase D (or AAVR2), which facilitates the transduction of both AAVR-dependent (Clade E AAVs) and-independent (AAV11 and AAV12) capsids^10–12^. Both AAVR and AAVR2 are Golgi-resident proteins comprising an ectodomain (ECD), a transmembrane domain (TM) and a carboxy-terminus tail (C-tail). In both receptors, the ECD binds AAV, while the C-tail facilitates intracellular trafficking^8,10,13^.

In this study, we have clarified the functional overlap between AAVR-and AAVR2-mediated transduction pathways and created a framework for designing receptor-guided platforms for vector testing. In defining the interactome of AAVR during productive AAV8 transduction, we discovered a direct physical interaction between AAVR and AAVR2, which mapped to non-AAV interacting C-terminal regions on both receptors. We then defined essential features within their C-tails and engineered chimeric AAV receptors with enhanced activity. Mechanistically, overexpression of AAVR or AAVR2 increased cell surface attachment and entry of AAV in a serotype-specific manner. Finally, we generated a stable cell line overexpressing a minimal functional AAVR2 (miniAAVR2) for enhanced *in vitro* transduction efficiency testing of AAVR2-engaging capsids including AAV8. These findings provide significant insights into AAV-receptor interactions which reinforce the need to optimise receptor-guided platforms to improve vector engineering.

## Results

### Proximity-dependent biotin identification (BioID) with AAVR identifies common viral receptor proteins

To identify proteins that interact with and potentially regulate AAVR function, we performed a BioID assay in human HuH-7 cells in the presence or absence of AAV8. BioID allows *in situ* labelling (biotinylation) of proximal proteins (<10 nM) within living cells^14,15^. First, we tested whether the fusion of BioID2 (a promiscuous biotin ligase mutant)^14^ at the C-terminus of AAVR impacts its receptor function. Lentiviral vectors containing mCherry were generated to express AAVR or an AAVR-BioID2 fusion with empty vector (EV) controls (**Fig. S1A**) and transduced in parallel into HEK293T *AAVR* KO cells. The overexpression of AAVR-BioID2 successfully rescued AAV8 transduction indicating that the fusion protein was functional as an AAV receptor (**Fig. S1B-C**).

After functional confirmation, we transduced HuH-7 cells with AAVR-BioID2-expressing lentiviruses (mCherry-P2A-AAVR-BioID2) or EV control (mCherry-P2A-BioID2). Cells positive for mCherry were enriched by fluorescence-activated cell sorting (FACS) followed by the addition of biotin for 16 h. In one experimental group, AAV8-eGFP was added to the cells for the last 30 min, whereas the other group did not receive AAV8. Cells were then lysed for analysis of total biotinylated proteins using affinity purification-mass spectrometry (AP-MS) or grown for a further 48 h to quantify AAV8 transduction (**Fig. 1A**). Robust expression of AAVR-BioID2-containing lentiviruses was confirmed in HuH-7 cells (>60%), based on mCherry fluorescence which occurred concomitantly with an increase in AAV8 transduction compared to control (**Fig. S1D**). Western blots were also performed with anti-AAVR antibody to demonstrate a modest overexpression of the AAVR-BioID2 fusion protein compared to endogenous AAVR and probed with streptavidin-HRP to confirm successful biotinylation (**Fig. 1B**).

**Figure 1:**
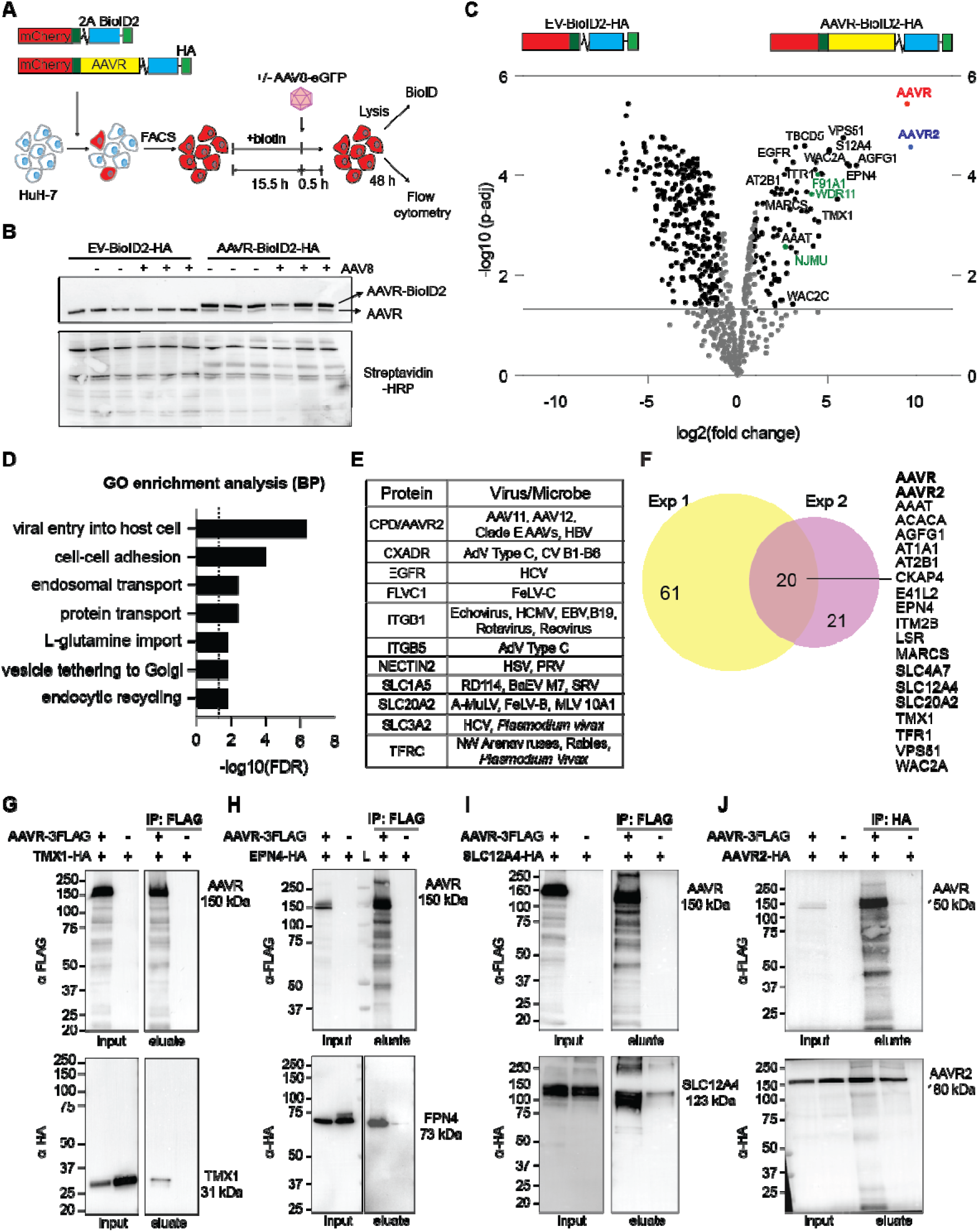
Proximity-dependent biotin identification (BioID) with AAVR identifies common viral receptor proteins. (A) BioID workflow used to identify AAVR-mediated protein-protein interactions during AAV8 transduction. HuH-7 cells were transduced with lentiviral vectors containing mCherry-2A-BioID2 (empty vector, EV) or mCherry-2A-AAVR-BioID2. Then, mCherry-positive cells were enriched by fluorescence-activated cell sorting (FACS). Biotin was added for 16 h, and AAV8-eGFP (5,000 vg/cell) was added 30 min before the completion of the experiment. Cells were either lysed for BioID and analysis with western blotting or grown for a further 48 h for flow cytometry analysis. (B) Western blot confirmation of AAVR-BioID2 expression in HuH-7 cells. Endogenous AAVR or ectopic AAVR-BioID2 expression was demonstrated using AAVR antibody (*upper panel*) and biotin labelling was confirmed using streptavidin-HRP (*lower panel*). (**C**) Representative volcano plot showing enriched proteins identified by AAVR BioID performed in HuH-7 cells in the presence of AAV8. AAVR or KIAA0319L (red), AAVR2 or CPD (blue) and WDR11 complex proteins (green) are highlighted. (**D**) Gene ontology (GO) enrichment analysis of 81 proteins identified in Exp 1 (log2(fold change)>1 and *p-adj*<0.05); BP=biological processes. (**E**) List of AAVR-interacting proteins which were previously confirmed to be receptors/entry factors: HBV=Duck Hepatitis B Virus; AdV=Adenovirus; CV=Coxsackievirus; HCV=Hepatitis C Virus; FeLV=Feline Leukemia Virus; HCMV=Human Cytomegalovirus; EBV=Epstein Barr Virus; B19=Parvovirus; HSV=Herpes Simplex Virus; PRV=Pseudorabies Virus; RD114=Feline Endogenous Retrovirus; BaEV M7=Baboon M7 Endogenous Virus; A-MuLV=Amphotropic Murine Leukemia Virus; *Plasmodium vivax*=malaria; NW=new world. (**F**) Venn diagram summarising two independent AAVR BioID experiments (Exp 1 and Exp 2). The proteins mutually enriched in both experiments are listed. (**G-J**) Confirmation of protein-protein interaction between AAVR and TMX1 (**G**), EPN4 (**H**), SLC12A4 (**I**) and AAVR2 (**J**) using co-immunoprecipitation (co-IP). AAVR-3FLAG as well as HA-tagged interactors were transfected into Expi293 cells and pulldown was performed using anti-FLAG beads (**G-I**) or anti-HA beads (**J**). Molecular weight markers are listed for each blot (kDa) along with the expected sizes of proteins and L=ladder in (**H**); (see also **Fig. S1** and **Supplementary data**).

Quantification of total biotinylated proteins (*p-adj*≤0.05 and log2FC≥1) showed a significant enrichment of several proteins implicated in AAV transduction, including AAVR2 (**Fig. 1C**). Gene ontology analysis revealed enrichment of biological processes which facilitate internalization of extracellular biologics comprising viral entry, endosomal transport, vesicle tethering to Golgi and endocytic recycling (**Fig. 1D**). Notably, various proteins previously identified as receptors for different classes of viruses were identified (**Fig. 1E**).

To verify these findings, we performed another independent screen, and 20 overlapping AAVR-interacting proteins were identified (**Fig. 1F**). Among the 20 overlapping candidates, were proteins with known or potential roles in AAV trafficking including transmembrane proteins (AAVR2, TFR1, EGFR and SLC12A4), Golgi-and retrograde transport-associated proteins (VPS51, AGFG1, TBCD5, EPN4/CLINT1 and TMX1) and proteins involved in nuclear transport and processing (WAC2A, CKAP4 and E41L2). Importantly, all three subunits of the WDR11 complex which facilitates the transport of acidic-cluster-containing proteins like AAVR and AAVR2 to the Golgi were also identified (WDR11, F91A1/FAM91A1 and NJMU/C17orf75)^16^. We also performed parallel BioID screens without the addition of AAV8 and observed fewer mutually enriched proteins (**Fig. S1E-F**). This suggests that ACACA, CKAP4, ITM2B, LSR, MARCS, SLC4A7, SLC12A4, E41L2 and WAC2A may only interact with AAVR in the presence of AAV8. Proteins identified in all four BioID screens are provided as **Supplementary data**.

To independently validate the BioID results we immunoprecipitated TMX1, EPN4 and SLC12A4, along with AAVR2. AAVR robustly immunoprecipitated: TMX1, an oxidoreductase which acts on transmembrane proteins (**Fig. 1G**); EPN4 or CLINT1, a clathrin-mediated trafficking protein (**Fig. 1H**); and SLC12A4, a cation-chloride transporter (**Fig. 1I**). AAVR2 efficiently co-immunoprecipitated AAVR (**Fig. 1J**). Collectively, these data identify and confirm the interactome of AAVR in the presence or absence of AAV8 and provide significant new insights into AAV receptor biology.

### AAVR and AAVR2 interaction is mediated by non-AAV binding residues

To map AAVR domains required for AAVR2 interaction (**Fig. 2A**), we tested a series of truncated AAVR constructs, including the N-terminal region (amino acids 1–311), PKD1–2 (AAV-binding polycystic kidney disease domains 1–2), PKD1–5 (all PKD domains 1–5), and the stalk region (amino acids 786-1049) comprising an α-helical segment, an EGF-like cysteine fold (C6), TM, and C-tail (**Fig. S2A**). Full-length AAVR was recovered following immunoprecipitation as expected, along with the AAVR stalk region with AAVR2 (**Fig. 2B**). This indicated that the AAVR stalk region is both sufficient and necessary for interaction with AAVR2. We then generated a series of AAVR2 ECD deletion mutants (ΔCP1, ΔCP2 and ΔCP3) to identify interacting domains within AAVR2 (**Fig. S2B**). Upon co-IP with AAVR2-HA, a reduction in interaction with full-length AAVR was observed with the ΔCP3 mutant (**Fig. S2C**). Additional mutants: ΔC-tail and ΔCP1-2 (**Fig. S2D**) were then tested via Co-IP with FLAG-tagged AAVR stalk. The AAVR stalk region could interact with ΔCP1-2 (containing CP3), but the interaction was lost upon deletion of CP3 alone (ΔCP3 mutant, **Fig. 2C**). These findings indicate CP3 is involved in AAVR interaction. However, the requirement for the AAVR2 C-tail is unclear as the ΔC-tail mutant was not detectable by HA antibody (as previously reported^10^).

**Fig. 2:**
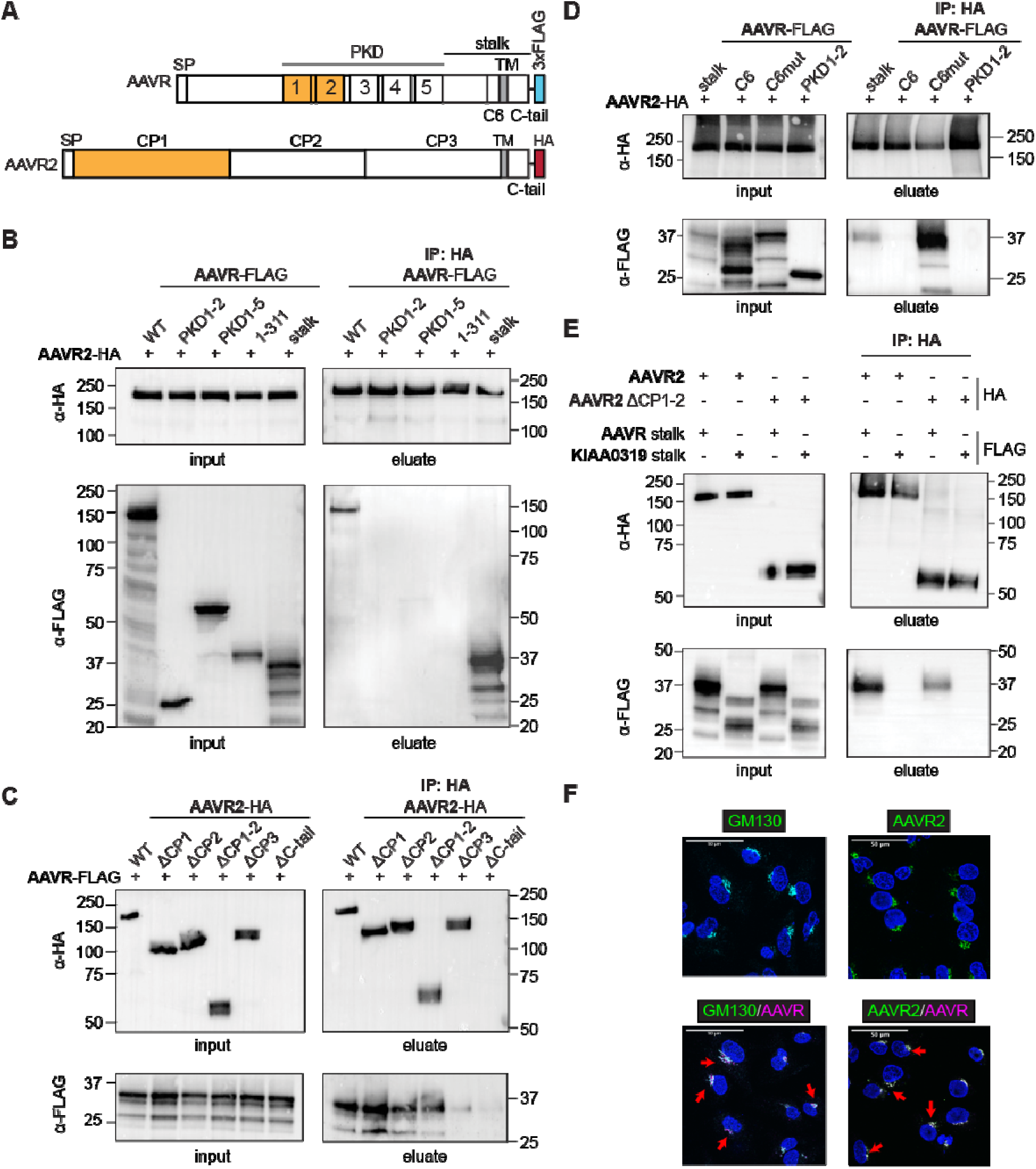
AAVR and AAVR2 interaction is mediated by non-AAV binding residues. (A) Schematic of full-length AAVR and AAVR2 constructs. Both proteins contain a signal peptide (SP), a transmembrane (TM) region, and a carboxy-terminal tail (C-tail). AAVR contains five polycystic kidney disease (PKD) domains in its ectodomain (ECD) and was tagged with a C-terminal 3xFLAG epitope. The stalk region contains an alpha-helical region, an EGF-like domain containing six cysteine residues (C6), a TM and C-tail. AAVR2 contains three carboxypeptidase-like domains (CP1, CP2 and CP3) in its ECD and was tagged with an HA epitope. Known AAV binding regions are coloured in orange. (**B**) Mapping of the minimal region in AAVR that interacts with AAVR2 by Co-IP. HA-tagged AAVR constructs included full-length AAVR (wildtype, WT), the N-terminal region (amino acid residues 1-311), the region containing all PKD domains (PKD1-5), the AAV interacting region (PKD1-2) and the stalk region. (**C**) Mapping of the minimal region in AAVR2 which interacts with the stalk region of AAVR by Co-IP. AAVR2 deletion mutants (Δ) were tested as follows: ΔCP1, ΔCP2, ΔCP1-2, ΔCP3 or ΔC-tail. (**D**) Refining the minimal AAVR interacting region with AAVR2 by Co-IP. AAVR deletion mutants included: stalk (positive control), C6 only (with C-tail), the stalk region with six cysteines mutated to alanine (C6mut) and PKD1-2 (negative control). (**E**) Determining the specificity of AAVR2 interaction with AAVR by Co-IP. WT or CP1-2-deleted-AAVR2 (ΔCP1-2) were used to pulldown AAVR stalk or the corresponding region in KIAA0319. (**B-E**) Schematics illustrating the regions in AAVR and AAVR2 used for Co-IP are shown in **Fig. S2A, B, D, E & G** respectively. Input and elution blots were loaded separately and each probed with anti-HA and anti-FLAG antibodies. Molecular weight markers (kDa) are indicated on all blots. (see also **Fig. S2**). (**F**) Immunofluorescence images showing the localization of AAVR, AAVR2 and GM130 (a Golgi marker) in HuH-7 cells. Merged panels show overlap within the Golgi with red arrowheads highlighting representative areas of colocalization; scale bar=50 μM.

To examine the juxtamembrane EGF-like C6 fold in AAVR of potential importance in protein-protein interactions^17^, we generated two AAVR stalk mutants: C6 (a stalk deletion mutant comprising only C6, TM and the C-tail); and a C6mut (full-length stalk with all 6 cysteines replaced by alanines) (**Fig. S2E**). Co-IP with AAVR2-HA showed a loss of AAVR2 interaction with C6, but not with the C6mut, further indicating the requirement of the entire AAVR stalk region for interaction (**Fig. 2D**). Protein alignment of the AAVR stalk and the stalk region of the closely related KIAA0319 protein revealed considerable similarity (66%) and common features including the EGF-like C6 domain (**Fig. S2F**). However, the KIAA0319 stalk region (**Fig. S2G)** was not pulled down by AAVR2-HA or the ΔCP1-2 mutant, demonstrating the specificity of the AAVR-AAVR2 interaction (**Fig. 2E**).

To determine the steady-state localization of AAVR and AAVR2, we performed immunofluorescence (IF) microscopy in HuH-7 cells. Confocal imaging revealed that both AAVR and AAVR2 show a predominantly perinuclear distribution pattern. Co-staining with the Golgi marker GM130 showed significant spatial overlap, confirming enrichment within the Golgi (**Fig. 2F**). Furthermore, a high degree of colocalization between AAVR and AAVR2 was also observed. Collectively, these results confirm a direct physical and spatial interaction between AAVR and AAVR2 in cells mediated by non-AAV binding residues located within their C-terminal regions.

### Trafficking signals within AAVR and AAVR2 C-tail regulate AAV uptake

Due to their critical roles in intracellular trafficking, we examined the molecular features of C-tails of AAVR and AAVR2. Their functional interchangeability was assessed by designing two chimeric receptors: AAVR with the C-tail of AAVR2 (AAVR-C2) and AAVR2 containing the C-tail of AAVR (AAVR2-C1). These HA-tagged chimeras were overexpressed in eHAP *AAVR* KO cells followed by transduction with different AAV serotypes (**Fig. 3A-C**). AAVR-C2 overexpression rescued the transduction of all tested serotypes at levels similar to wild type (WT) AAVR. AAVR2 incorporating the AAVR C-tail (AAVR2-C1) selectively rescued the transduction of Clade E AAVs (AAV8, AAVrh10 and AAVhu37), similar to WT AAVR2 (**Fig. 3D**). These results demonstrate that the C-tails are functionally interchangeable between the two AAV receptors and offer the possibility of designing chimeric receptors with altered functionality.

**Fig. 3:**
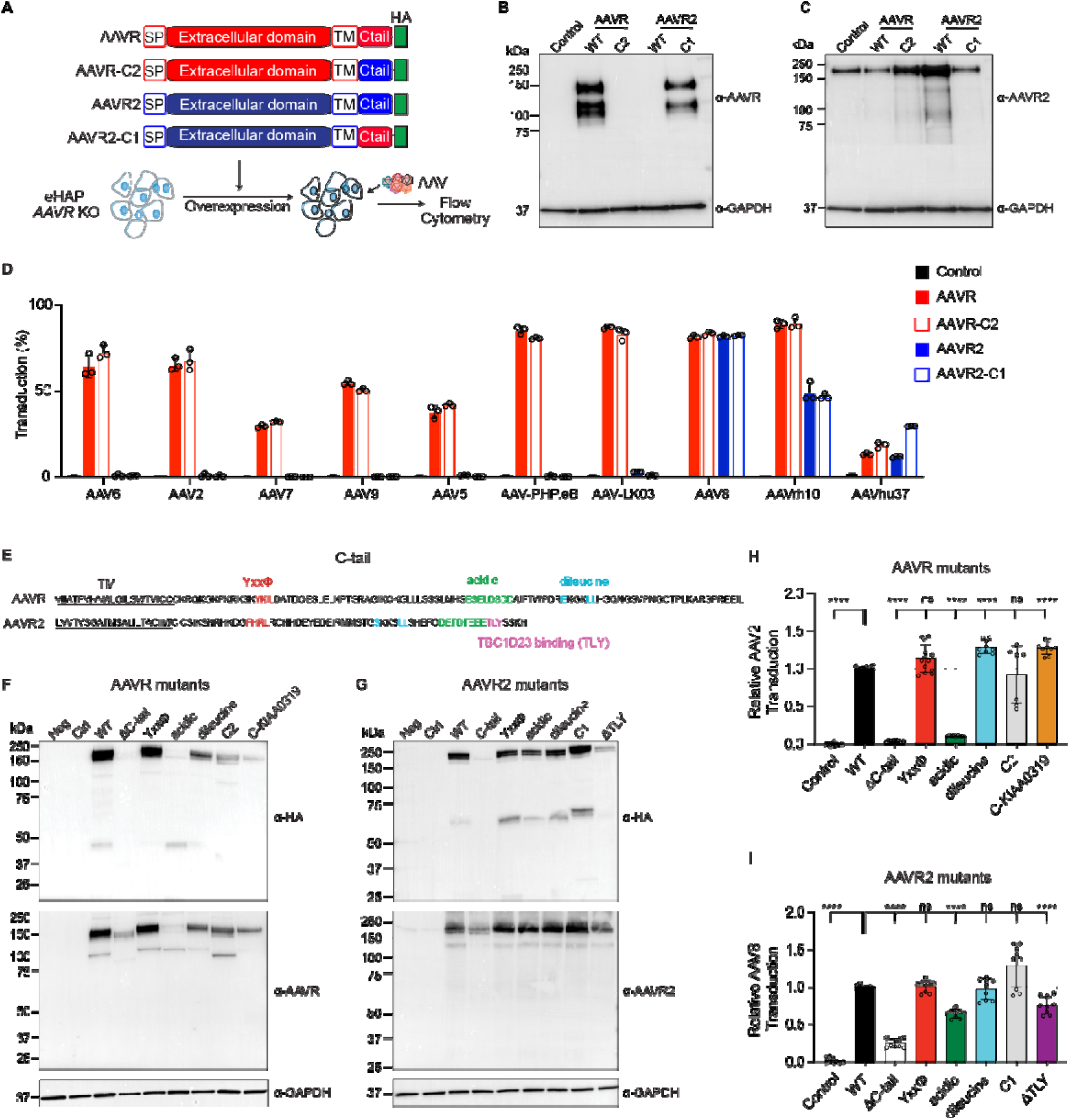
Trafficking signals within AAVR and AAVR2 C-tails regulate AAV uptake. (**A**) Strategy to test the functional interchangeability of AAVR and AAVR2 C-tails. AAVR with AAVR2 C-tail (AAVR-C2) and AAVR2 with the C-tail of AAVR (AAVR2-C1) were overexpressed using lentiviruses in eHAP *AAVR* KO cells along with wild-type (WT) AAVR or AAVR2 followed by transduction with a range of AAV serotypes. (**B-C**) Representative western blots performed with anti-AAVR (**B**) or anti-AAVR2 (**C**) antibodies in eHAP *AAVR* KO cells overexpressing AAVR WT or C2, or AAVR2 WT or C1. GAPDH was used as a loading control. (**D**) Transduction of different AAV serotypes (100,000 vg/cell) in eHAP *AAVR* KO cells overexpressing AAVR, AAVR-C2, AAVR2 or AAVR2-C1 (*n*=3). (**E**) Schematic of amino acid sequences of known and putative trafficking motifs in the AAVR and AAVR2 C-tails. (**F-G**) Representative western blots performed with anti-HA, anti-AAVR, anti-AAVR2 or anti-GAPDH antibodies to confirm the expression of different C-tail mutants of AAVR and AAVR2. (**H-I**) Transduction of AAV2-eGFP (10,000 vg/cell) in HEK293T *AAVR* KO cells transfected with expression vectors to overexpress WT AAVR or AAVR C-tail mutants (**H**), and WT AAVR2 or AAVR2 C-tail mutants (**I**), measured using flow cytometry. (**H & I**) Data represent mean ± SD of three or four independent experiments each performed in triplicate. Transduction in mutant-overexpressing cells was normalized to that in WT AAVR-or AAVR2-overexpressing cells and empty vector transfected cells were used as controls. All statistical analysis were performed using the Welch’s t-test, where \**p*<0.05, \*\**p*<0.01, \*\*\**p*<0.001, \*\*\*\**p*<0.0001 and ns=not significant.

Analysis of AAVR and AAVR2 from publicly available proteomics data showed extensive post-translational modifications (PTMs) within the C-tails of both receptors compared to the ectodomains of both proteins (**Fig. S3A-B**). We also identified motifs implicated in protein sorting and trafficking including the canonical sorting motif YXXΦ in AAVR (YKIL) and a variant of this in AAVR2 (FHRL), acidic clusters and dileucine motifs^18,19^ (**Fig. 3E**). To determine the functional relevance of these motifs in AAV entry, we generated a panel of C-tail mutants. For AAVR, the following C-tail mutants were generated: complete C-tail deletion (ΔC-tail), YXXΦ mutation, acidic cluster mutation, dileucine (ExxxLL) mutation, AAVR-C2 and a chimeric AAVR with the C-tail of the related KIAA0319 protein. For AAVR2, mutants included ΔC-tail deletion, FHRL mutation, acidic cluster mutation, dileucine mutation and AAVR2-C1 chimera. In addition, we examined the mutation of the TBC1D23-binding motif (TLY)^20^.

All mutants were transfected into HEK293T *AAVR* KO cells followed by transduction with AAV2 (for AAVR mutants) or AAV8 (for AAVR2 mutants) (**Fig. 3F-G**). The removal of the C-tail, or the acidic cluster within the C-tail of AAVR, significantly inhibited its ability to rescue AAV2 transduction in *AAVR* KO cells (*p*<0.0001, **Fig. 3H**). AAVR incorporating the KIAA0319 C-tail rescued AAV2 at a level higher than WT AAVR (*p*<0.0001), despite lower expression by western blot. Similarly, the ΔC-tail and acidic cluster mutants of AAVR2 rescued the transduction of AAV8 at lower levels compared to WT AAVR2 (*p*<0.0001, **Fig. 3I**). The AAVR2 TLY mutant showed reduced activity indicating a potential role of TBC1D23 binding in AAV uptake. Together, these results identify critical C-tail motifs in AAVR and AAVR2 required for AAV cellular entry.

### C-tail optimization enhances AAVR2 activity

HA-tagged chimeric expression constructs were generated by fusing the TM and C-tails of AAVR or AAVR2 with the ECD of CD8 as a cell surface reporter. These constructs were transfected into HEK293T cells followed by flow cytometric quantification of CD8 cell surface expression and transfection efficiency (HA expression). Increased CD8 surface expression at comparable levels of HA expression indicates that the C-tail enhances cell surface trafficking.

The cell surface expression of CD8 fused to the TM and C-tails of archetypal trafficking proteins AAVR, AAVR2, CI-MPR and FURIN along with WT CD8 was measured (**Fig. 4A**). CD8-C-tail chimeras were transfected into HEK293T cells, and expression was quantified using flow cytometry (**Fig. 4B**) and western blotting (**Fig. 4C**). All C-terminal tails reduced CD8 surface expression relative to WT CD8; however, the AAVR2 C-terminal tail caused the greatest reduction, with significantly lower expression than all other C-tails tested (**Fig. 4D-E**).

**Fig. 4:**
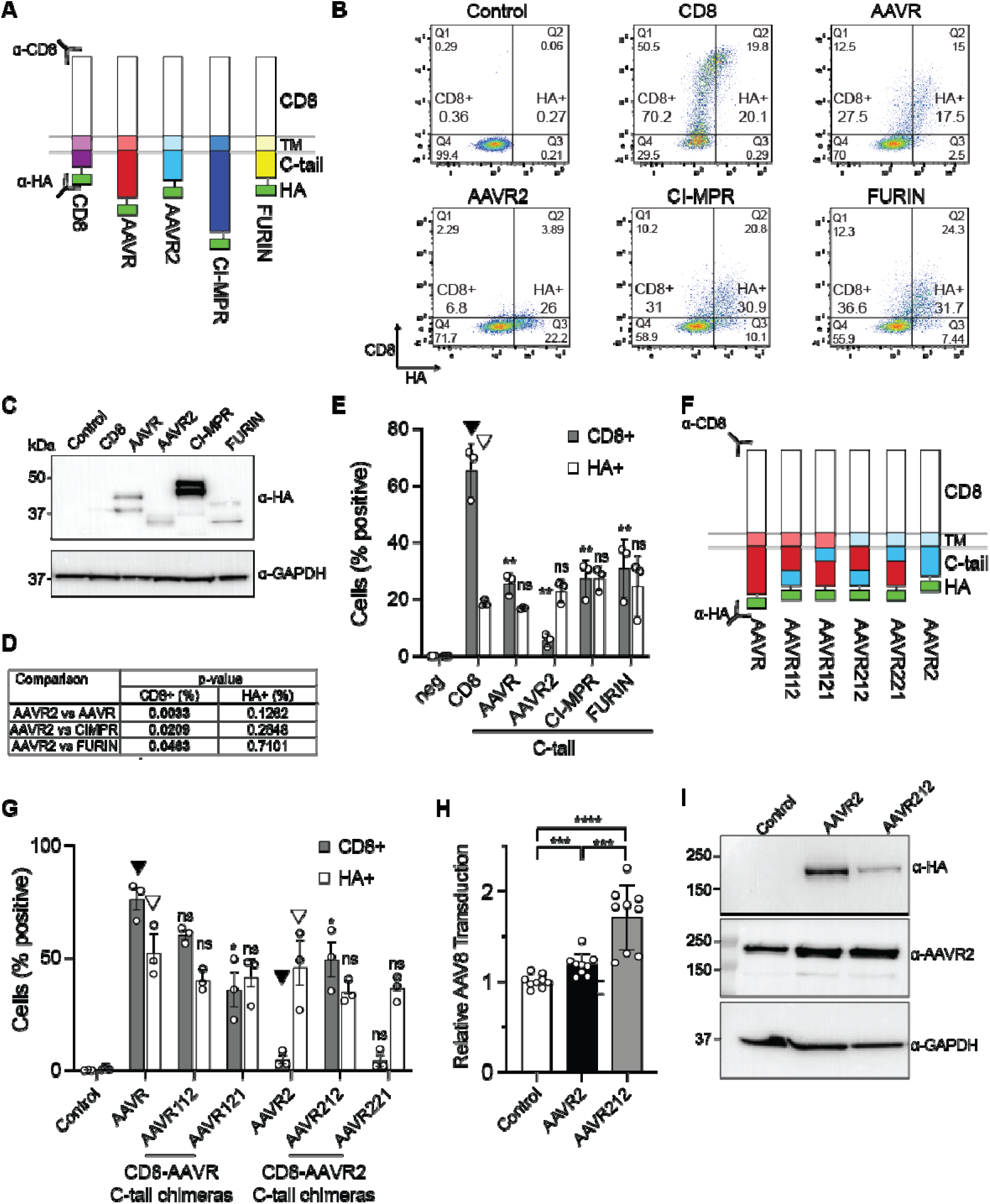
C-tail optimization enhances AAVR2 activity. **(A**) Schematic of CD8 fusion reporters consisting of the CD8 ectodomain with the TM and C-tail of archetypal trafficking proteins AAVR, AAVR2, CI-MPR and FURIN. The CD8 TM with C-tail was used as control. Antibodies against CD8 and HA (depicted) were used to determine cell surface staining and total protein expression respectively. (**B-C**) HEK293T cells were transfected with CD8-based chimeric constructs for 48 h before lysis for Western blotting, or staining fresh cells with CD8 antibody before fixing, permeabilization and staining with HA antibody, and performing flow cytometric analysis.(**B**); or Western blot analysis (**C**). (**D**) Statistical analysis comparing AAVR2 C-tail trafficking with other C-tails. (**E**) Summary of flow cytometric analysis of CD8 and HA expression depicting percentage of cells with CD8 surface expression and total HA expression. Arrowheads indicate the sample used for each statistical comparison for CD8+ and HA+ samples. (F) Schematic of CD8 fusion reporter constructs with different chimeric AAVR and AAVR2 C-tails. (G) Summary of flow cytometric analysis of CD8 and HA expression depicting percentage of cells with CD8 surface expression and total HA expression. Arrowheads indicate the sample used for each statistical comparison for CD8+ and HA+ samples. For all CD8 reporter assays, *n*=3 independent experiments performed in triplicate and data represent mean ± SD with statistical analysis determined with a Welch’s t-test where \**p*<0.05 and \*\**p*<0.01. (**H**) AAV8 transduction (10,000 vg/cell) in HEK293T *AAVR* KO cells overexpressing WT AAVR2 or AAVR212. (**I**) Representative western blots performed with antibodies against HA, AAVR2 and GAPDH in HEK293T *AAVR* KO cells overexpressing AAVR2 or AAVR212; (see also **Fig. S4**).

To optimize the AAVR2 C-tail for enhanced trafficking, chimeric C-tails were generated by domain swapping with the AAVR C-tail (**Fig. 4F**). We generated AAVR2 C-tail chimeras AAVR212 and AAVR221, while the AAVR C-tail chimeras AAVR112 and AAVR121 were generated for comparison. These C-tails were fused to the CD8 ECD and used for the CD8 trafficking assay in HEK293T cells (**Fig. S4A-B**). When compared to the WT AAVR C-tail, a significant decrease in CD8 surface expression was observed for the AAVR121 chimera (*p*<0.05), indicating the major role of residues 954-1000 in AAVR-mediated trafficking (**Fig. 4G, Fig. S4A**). In contrast, the AAVR212 chimera led to significantly higher CD8 surface expression compared to the WT AAVR2 C-tail (10.3-fold, *p*<0.05, **Fig. 4G, Fig. S4B**).

To further refine the C-tails, we generated additional C-tail chimeras for AAVR (AAVR1J21 and AAVR1Y21) and AAVR2 (AAVR2J12 and AAVR2Y12) (**Fig. S4C**). These chimeras were fused to the CD8 ECD and transfected into HEK293T (**Fig. S4D**). As expected, compared to the WT AAVR C-tail, AAVR121 showed a significant reduction of CD8 surface expression (*p*<0.05) but not AAVR1J21 and AAVR1Y21 (**Fig. S4E-F**). The AAVR2 C-tail chimeras AAVR2J12 and AAVR2Y12 supported a significantly higher surface expression of CD8 compared to the WT AAVR2 C-tail (*p*<0.05), however, AAVR212 was superior (**Fig. S4E-F**). We also compared AAVR212 with the AAVR2K2 mutant containing the corresponding C-tail region from KIAA0319 instead of AAVR (**Fig. S4G-H**). AAVR212 supported significantly higher surface expression of CD8 compared to AAVR2K2 indicating a higher trafficking ability of these residues in AAVR compared to KIAA0319 (**Fig. S4I**).

A full-length AAVR212 and WT AAVR2 were compared in their ability to rescue AAV8 transduction in HEK293T *AAVR* KO cells. AAVR212 overexpression led to significantly higher AAV8 transduction compared to WT AAVR2 ((*p*<0.001), **Fig. 4H**) despite a lower level of protein expression determined by western blot (**Fig. 4I**). When combined, these results identify key functional residues within the AAVR and AAVR2 C-tails required for efficient trafficking and prompts the rational design of AAV receptors with enhanced functions.

### Receptor overexpression enhances *in vitro* AAV transduction assays

Finally, we focused on the mechanisms and potential applications of receptor overexpression. We previously showed that AAV8 transduction is enhanced following the overexpression of AAVR or AAVR2. Here, we assessed the dose-dependent effects of AAVR and AAVR2 on AAV8 transduction. HuH7 cells were transduced with different volumes of lentiviruses expressing AAVR or AAVR2 (**Fig. S5A-B**), followed by AAV8-eGFP. As shown in **Fig. S5C-D**, AAV8 transduction increased in a dose-dependent manner upon overexpression of AAVR or AAVR2. We next quantified the cell surface expression of AAVR and AAVR2 using flow cytometry at the highest dose (400 µL). There was a significant increase in the cell surface expression of AAVR (*p*<0.05, **Fig. S5E-F**) and AAVR2 (*p*<0.05, **Fig. S5G-H**) compared to control indicating that an increase in the surface expression of the receptors correlates with enhanced AAV8 transduction.

To examine cellular attachment and entry, AAV8 was added to eHAP cells overexpressing AAVR or AAVR2 followed by an incubation at 4°C for 1 h (attachment) or 37°C for 3 h (entry) (**Fig. 5A**). As shown in **Fig. 5B-C**, AAV8 staining was below the detection limit in control cells consistent with inefficient AAV8 cellular entry^11^. Upon overexpression of AAVR or AAVR2, markedly increased staining for AAV8 was observed in both attachment and entry conditions. For both conditions, we observed a significantly higher colocalization of AAV8 with AAVR or AAVR2 in overexpression groups compared to controls (**Fig. 5D**). Additionally, we observed a linear trend consistent with a dose-response relationship between AAV8 and AAVR or AAVR2 in both attachment and entry conditions (**Fig. S5I-J**). These results indicate that AAVR or AAVR2 are dose-limiting for cellular attachment and entry of AAV8.

**Figure 5:**
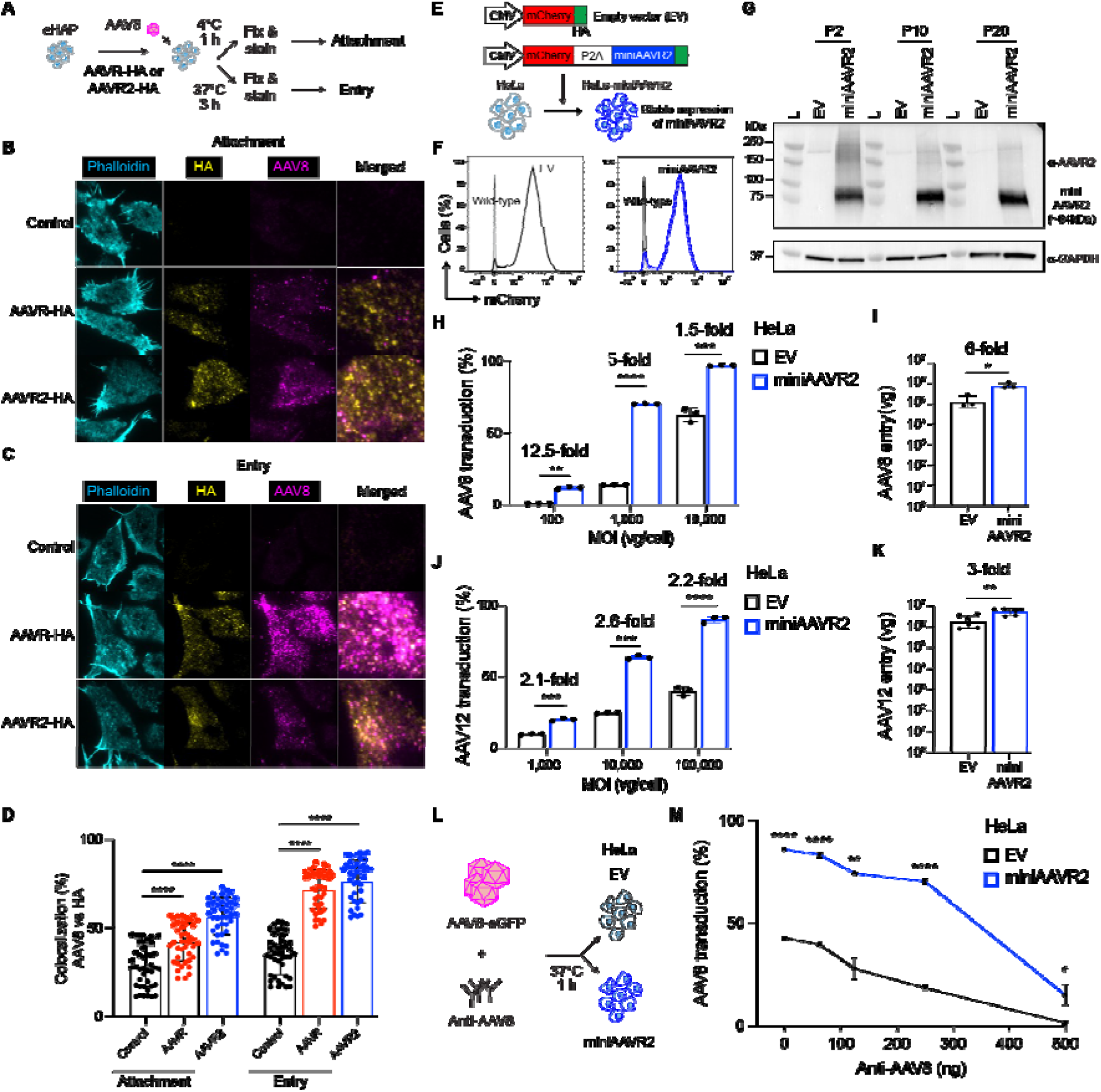
Receptor overexpression enhances *in vitro* transduction assays. (**A**) Schematic of immunofluorescence-based AAV8 cellular attachment and entry assays performed in eHAP cells. HA-tagged AAVR or AAVR2 were overexpressed using lentiviral vectors for 48 h followed by the addition of AAV8 (100,000 vg/cell). Cells were incubated at 4°C for 1 h (attachment) or 37°C for 3 h (entry), fixed and stained with phalloidin or antibodies against HA or AAV8. (**B-C**) Representative immunofluorescence images performed with phalloidin (blue), anti-HA (representing AAVR or AAVR2) (yellow) and anti-AAV8 (magenta) antibodies in control eHAP cells or cells overexpressing AAVR or AAVR2 at attachment (**B**) and entry (**C**) conditions. (**D**) The percentage of AAV8 and HA co-localization signals in eHAP control or AAVR-and AAVR2-overexpressing cells. Signal intensities of AAV8 and HA in different cells are shown (n=42 each bar). (**E**) Generation of HeLa cells stably expressing miniAAVR2 (HeLa-miniAAVR2). Cells were transduced with a lentiviral vector expressing mCherry and miniAAVR2 separated by a self-cleaving P2A peptide driven by the ubiquitous cytomegalovirus (CMV) promoter versus empty vector (EV) control. (**F**) Overlay plots representing mCherry expression in wild type vs EV transduced or HeLa-miniAAVR2 cells (x-axis=mCherry and y-axis=percentage of cells at passage 20). (**G**) Western blot performed with anti-AAVR2 and anti-GAPDH on lysates from HeLa-EV or - miniAAVR2 cells at indicated passage (P) numbers. (**H**) Transduction of AAV8-eGFP in HeLa-EV or-miniAAVR2 cells at indicated multiplicity of infection (MOI) measured as vector genomes/cell (vg/cell). (**I**) Total number of AAV8-eGFP genomes detected in HeLa-EV or-miniAAVR2 cells 24 h post-administration s (MOI=100,000 vg/cell). (**J**) Transduction of AAV12-eGFP in HeLa-EV or - miniAAVR2 cells at indicated MOI. (**K**) Total number of AAV12-eGFP genomes in HeLa-EV or - miniAAVR2 cells (MOI=100,000 vg/cell). (**L**) Schematic of neutralizing antibody (NAb) assay performed with anti-AAV8 antibody in HeLa-EV or-miniAAVR2 cells. (**K**) Transduction of AAV8-eGFP in HeLa-EV or-miniAAVR2 cells after being incubated with the indicated amount of anti-AAV8 antibody. (**H-M)** *n*=3, Welch’s t-test \**p*<0.05, \*\**p*<0.01, \*\*\**p*<0.001 and \*\*\*\**p*<0.0001. (see also **Fig. S5**).

We had previously defined the minimal functional AAVR2 (miniAAVR2) consisting of SP, CP1, TM and C-tail^10^. To enable standardized measurement of transduction of AAVR2-engaging gene therapy vectors, we generated HeLa cells stably expressing miniAAVR2 (HeLa-miniAAVR2) or EV control (HeLa-EV). HeLa cells were transduced with lentiviral vectors co-expressing mCherry and miniAAVR2 and cells with high mCherry expression were enriched with FACS (**Fig. 5E-F**). Stable expression of miniAAVR2 was maintained in HeLa-miniAAVR2 cells up to and including the highest passage number tested (Passage 20, **Fig. 5G**). There was an increase of 12.5-, 5-and 1.5-fold in AAV8 transduction at MOIs of 100, 1,000 and 10,000 vg/cell respectively (**Fig. 5H**), accompanied by a 6-fold increase in cellular entry for AAV8 (**Fig. 5I**) in HeLa-miniAAVR2 cells compared to EV control. Similarly, AAV12 transduction increased by 2.1-, 2.6-and 2.2-fold at 1,000, 10,000 and 100,000 vg/cell respectively (**Fig. 5J**) with a 3-fold increase in cellular uptake (**Fig. 5K**).

Next, we tested whether HeLa-miniAAVR2 cells exhibit enhanced sensitivity for detecting antibody-mediated neutralization of AAV8 (**Fig. 5L**). AAV8-eGFP was pre-incubated with increasing amounts of a monoclonal antibody against AAV8 and used to transduce EV control or HeLa-miniAAVR2 cells. HeLa-miniAAVR2 cells showed significantly higher levels of transduction across all antibody concentrations tested (**Fig. 5M**, *p*<0.05). Notably, at an antibody amount of 500 ng, complete AAV8 neutralization was seen in control cells but remained detectable in HeLa-miniAAVR2, indicating increased sensitivity (**Fig. 5M)**.

To further establish the applications of miniAAVR2-based cell systems, we overexpressed miniAAVR2 in a panel of cell lines and observed a significant increase in AAV8 transduction in all common cell adherent and suspension cell lines tested compared to controls (*p*<0.05, **Fig. S5K**). This indicates that miniAAVR2 overexpression can be adapted to define AAV8 transduction in a diverse range of cells. Taken together, these results indicate that overexpressing AAVR or AAVR2 enhances surface attachment and entry of AAV8 and supports the potential of receptor-based cell systems as improved tools for AAV vector development.

## Discussion

Our discovery of AAVR2 (Carboxypeptidase D) as an alternative AAV receptor significantly refined the biological understanding of AAV entry. AAVR2 not only influences the transduction of exclusively AAVR2-dependent serotypes (AAV11 and AAV12) but also facilitates the transduction of clinically relevant AAVR-dependent Clade E AAVs (including AAV8)^10^, highlighting a previously unrecognized complexity in receptor usage. In the current study, we elucidate the mechanistic interplay between AAVR and AAVR2, identify functionally important components of both receptors and provide a framework for receptor-guided generation of tools to assist the development of AAV-based gene therapies.

Our delineation of the AAVR interactome with or without AAV8 reveals previously unknown AAV receptor biology (**Fig. 1**). AAVR BioID experiments identified several host factors directly regulating AAV transduction like AAVR2^10^, VPS51^9^ and TFR1^21^ or protein complexes involved in intracellular trafficking of AAV, such as the members of the solute carrier family^8,22,23^, Golgi-associated retrograde protein (GARP) complex proteins and the Wiskott-Aldrich Syndrome protein and SCAR Homologue complex (WASH) complex proteins^9^. Future studies examining the serotype specificity of these host proteins will further increase our understanding of AAV transduction biology.

We show that the interaction between AAVR and AAVR2 is mediated by non-AAV-binding residues located in the C-terminal region of both receptors (**Fig. 2**). Furthermore, this interaction was also observed in the absence of AAV, revealing that AAVR-AAVR2 association can occur independently of AAV binding. The co-location of both receptors within the Golgi strengthens our previous understanding that this organelle serves as a critical convergence point and intracellular depot for AAV trafficking^7^. This colocalization also supports an intracellular trafficking model where diverse AAV serotypes, despite utilizing different primary entry factors, share common transport machineries. Further studies examining this interaction in the context of transduction of different serotypes will be of significant interest.

Our observation that the AAVR2 C-tail is the least efficient in supporting trafficking among the 5 archetypal trafficking proteins examined, provided the rationale for sequence optimization to enhance trafficking activity. We functionally characterized trafficking signals in AAVR and AAVR2 C-tails and designed several chimeras with enhanced functions (**Fig. 3**). We were also able to significantly enhance the AAVR2 C-tail’s ability to support AAV8 transduction by domain swapping (**Fig. 4)**.

In the context of receptor expression, we previously reported that the cell surface attachment and entry of AAVR2-dependent capsids like AAV11 are not reduced despite a loss of overall transduction in *AAVR2* KO cells^10^. Similarly, others have shown that *AAVR* KO does not impact the surface attachment and entry of AAVR-dependent capsids but affects intracellular trafficking upstream to nuclear entry^12,24^. We therefore sought to define the extent to which receptor overexpression increases overall transduction. Using imaging-and qPCR-based assays we observed an increased AAV8 cellular attachment and entry in AAVR-or AAVR2-overexpressing cells (**Fig. 5**). The increased surface expression of the receptors in these cells indicate that the first contact between AAV-AAVR or AAV-AAVR2 may occur at the cell surface but is limited or transient due to the limited availability of the receptors. Due to the inefficient nature of AAV8 cellular entry and unavailability of reliable antibodies against AAV11 and AAV12, we were unable to perform a similar immunofluorescence-based study in *AAVR* and *AAVR2* KO cells at steady-state. However, upon receptor overexpression, AAV8 signals were readily detected and dispersed throughout the cells, indicating that at least a fraction of AAV particles may remain associated with receptors post-Golgi processing.

To enhance *in vitro* transduction testing of AAVR2-engaging serotypes, we generated a cell line stably expressing a minimal functional AAVR2 with enhanced permissivity. We observed an increased sensitivity in the detection of antibody-mediated neutralization of AAV8 in these cells. Thus, this cell line could be a valuable tool for testing the potency of clinically important Clade E AAVs which transduce cell lines poorly. This was further demonstrated by our miniAAVR2 overexpression data which confirmed increased AAV8 transduction in ten different cell lines.

Our study provides significant insights into the biology of AAV receptors and identifies potential mechanisms driving the serotype-specific nature of AAV tropism. Further, our results provide evidence that AAV receptor engineering could complement capsid optimization as a valuable tool in the field of AAV vectorology and gene therapy.

## Materials and methods

### Cell lines

All cell lines except HuH-7 (CellBank Australia) and eHAP (Horizon Discovery, CO, USA) were obtained from ATCC (American Type Culture Collection, VA, USA). HEK293T, Calu-6, CHO-K1, HaCaT, Hep3B, HuH-7, HFF, Mero-95, MIA PaCa-2, and U-87 MG cell lines were grown in DMEM (Gibco, MA, USA; 11965175) supplemented with 10% (v/v) FBS (Bovogen, VIC, Australia; SFBS) and 1% (v/v) penicillin/streptomycin (Gibco; 15140122). eHAP cells were grown in IMDM (Gibco; 12440053) supplemented with 10% (v/v) FBS and 1% (v/v) penicillin/streptomycin and the THP-1 cell line was maintained in RPMI 1640 (Gibco; 11875093) supplemented with 10% (v/v) FBS and 1% (v/v) penicillin/streptomycin.

### AAVR proximity-dependent biotin identification (BioID)

A promiscuous biotin ligase variant, BioID2^14^, was obtained from plasmid MCS-13X-Linker-BioID2-HA (Addgene #80899). Lentiviral plasmids were constructed to generate pFUW-mCherry-2A-2xGS-BioID2-HA (empty vector; EV-BioID2-HA) and pFUW-mCherry-2A-AAVR-2xGS-BioID2-HA (AAVR-BioID2-HA). VSV-G–pseudotyped lentiviral supernatants were produced and used to transduce HuH-7 cells. Cells positive for mCherry were enriched by FACS and expanded for BioID analysis. For each cell population, six 10 cm dishes (6×10^6^ cells/dish) were seeded, and at ∼80% confluence, cells were treated with 50 µM biotin (Sigma, MO, USA; B4639) overnight. After 15.5 h, AAV8-eGFP (5,000 vg/cell) was added to three dishes per condition and incubated for 30 min at 37°C. Most cells were then harvested, washed four times in cold PBS (300 g), and pelleted, while a subset was maintained in culture (with virus removed) for an additional 48 h for flow cytometric analysis. Each cell pellet was lysed in 550 µL lysis buffer (50 mM Tris-HCl pH 7.5, 150 mM NaCl, 1% (w/v) n-dodecyl-β-D-maltoside, 0.5% (w/v) digitonin, 0.5% (w/v) sodium deoxycholate, 1 mM EDTA, 1 mM EGTA, 1 mM PMSF, 1 mM DTT, 1× protease inhibitor cocktail (cOmplete, Roche Life Sciences, Mannheim, Germany; 4693132001), 2 U/mL Universal Nuclease (ThermoFisher, MA, USA; 88700). Lysates were vortexed and sonicated on ice (5 cycles, 1 min on/10 s off) until clear, then centrifuged at 20,000 g for 20 min. Clarified supernatants were collected, with 20 µL reserved as input, and the remainder incubated with 40 µL streptavidin–agarose bead slurry (Merck, MA, USA; GE28-9857-99) overnight at 4 °C with rotation. Beads were centrifuged at 800 g for 5 min, supernatant was removed, and beads were washed four times with 800 µL Wash Buffer I (50 mM Tris-HCl pH 7.5, 150 mM NaCl), centrifuging at 800 g between washes. For on-bead digestion, beads were resuspended in Digestion Buffer I (2 M urea, 50 mM Tris-HCl pH 7.5, 1 mM DTT, 3 µg trypsin (Promega, WI, USA; V5117), 1 µg LysC (Promega; VA1170) and incubated at 30°C, 1,200 rpm for 2 h. Beads were then centrifuged at 800 g, and the supernatant transferred to a LoBind tube (Eppendorf, Hamburg, Germany; 0030108116). Beads were resuspended in an equal volume of Digestion Buffer II (2 M urea, 50 mM Tris-HCl pH 7.5, 5 mM iodoacetamide), vortexed, centrifuged at 2,000 g, and this supernatant was combined with the first. Digestion was quenched with formic acid to a final concentration of 2% (v/v). Peptides were desalted using C18 tips (Millipore, MA, USA; ZTC18S960). Tips were conditioned with 70 µL 80% (v/v) acetonitrile/5% (v/v) formic acid and centrifuged at 1,000 g for 1 min, then equilibrated twice with 70 µL 5% (v/v) formic acid (1,000 g, 1 min each). Samples were then loaded onto the tips and centrifuged at 1,000 g for 1 min. Tips were washed four times with 70 µL 0.1% (v/v) formic acid (1,000 g, 1 min). Peptides were eluted into new LoBind tubes with 10 µL 50% (v/v) acetonitrile/0.1% (v/v) formic acid, centrifuged at 1,000 g for 3 min, and elution was repeated once. Eluates were dried in a SpeedVac and stored for mass spectrometry.

### Mass spectrometry

Raw mass spectrometry data were generated using the Q Exactive HF-X Quadrupole-Orbitrap system and analysed as described in our previous work^25,26^. In brief, data were processed using MaxQuant (version 2.7.5.0) with standard parameters and label-free quantification (LFQ) enabled^27^. Carbamidomethylation of cysteine was set as a fixed modification and methionine oxidation as a variable modification. LysC/trypsin were specified as proteolytic enzymes, allowing up to two missed cleavages to account for lysine biotinylation, and a 1% false discovery rate was applied at both peptide and protein levels. LFQ values were analysed in R using LFQ-Analyst with missing values imputed using the Perseus algorithm^28^. Proteins with log□fold change>1 and *p-adj* <0.05 were considered significantly enriched.

### Western Blotting

Cells were harvested in a lysis buffer containing Tris-HCl (20 mM), NaCl (150 mM), Triton X-100 (1% v/v), SDS (0.1% v/v), sodium deoxycholate (0.5% w/v) and protease inhibitor cocktail. Electrophoresis was performed on SDS-NuPAGE gels followed by transfer onto PVDF membranes. The membranes were blocked with skim milk (5% w/v) in PBS containing Tween 20 (0.1% v/v, PBST). and probed with antibodies against AAVR (1:5,000; Abcam, Cambridge, UK; ab105385), AAVR2 (1:5,000; Abcam; ab184960), GAPDH (1:5,000; Abcam; ab8245), FLAG (1:5,000; Sigma-Aldrich, A8592-1) or HA (1:5,000; Biolegend, CA, USA; 901501). After washing with PBST, membranes were probed with appropriate horseradish peroxidase (HRP)-conjugated secondary antibodies. The ECL western blotting substrate (ThermoFisher; 32209) was used to develop the blots, and visualization was performed using a Bio-Rad Chemidoc imaging system.

### Co-immunoprecipitation (Co-IP)

Plasmids containing full-length AAVR (pCMV-AAVR-3xFLAG) and AAVR2 (pCMV-AAVR2-HA) were used to make additional deletion or point mutants for Co-IP experiments. PCR primers used to generate AAVR or AAVR2 domain mutants are listed in **Table S1**. Plasmids expressing potential interactors included TMX1 (pCMV6-TMX1-eGFP; Origene, MD, USA; RC206187L3), CLINT1/EPN4 (Horizon Discovery; BC004467), and SLC12A4, which were used to make pcDNA3.1-based HA-tagged constructs for CoIP validation. For Co-IP, Expi293 cells (3×10^6^) were transfected with 2 μg of each plasmid DNA (4 μg total) using polyethylenimine (Polysciences, PA, USA; PEI). Cells were incubated for 48 h with shaking, collected, and washed twice with PBS. Cells were lysed in 600 μL TBS buffer (50 mM Tris-HCl, 200 mM NaCl, 0.1% (v/v) Triton-X-100) containing freshly added protease inhibitor cocktail, 1 mM PMSF (Sigma, 10837091001), 0.2 mM DTT, 0.2% (w/v) DDM, 0.2% (w/v) digitonin and 1 mM EDTA. Lysates were sonicated for 8 cycles of 30 s on/ 30 s off in low power mode and clarified by centrifugation at 16,000 g for 20 min at 4 °C. Supernatants were transferred to a new tube and an input sample (40 μL) was reserved, to which 15 μL 4x LDS with 10% (v/v) β-mercaptoethanol was added and heated for 15 min at 80 °C. For each pulldown, 35 μL of anti-HA agarose bead slurry (ThermoFisher, 26182) or anti-FLAG agarose bead slurry (Merck; A2220) was used. Cell lysate (∼560 μL) was added to the beads and rotated overnight at 4 °C. The beads were centrifuged for 5 min at 1,000 g (4 °C) and the supernatant was removed. The beads were carefully washed 5 times in 1 mL of Wash Buffer 1 (50 mM Tris pH 7.5, 200 mM NaCl, 0.25% (v/v) IGEPAL) with rotation for 5 min at 4 °C before centrifugation at 1,000 g. Wash Buffer 2 (10 mM Tris-HCl pH 7.5, 100 mM NaCl, 0.1 mM EDTA) was then added (1 mL) and the beads rotated for 3 min at 4 °C before centrifugation at 1,000 g. To elute proteins, 25 μL of HA peptide (200 μg/mL, Merck; 11249) or FLAG peptide (150 μg/mL, Merck; F4799) was added to the beads and incubated at 4 °C on a shaker for 40 min at 1,200 rpm. The beads were spun at 1,000 g for 2 min, and the supernatant transferred to a new tube and the elution was repeated once more. TBS (10 μL) was mixed with the beads and centrifuged, combined with the other eluates (60 μL total), and then 4xLDS with 10% (v/v) β-mercaptoethanol was added. Input (10 μL) and eluate samples (35 μL) were subjected to SDS-PAGE.

### CD8-based cell surface trafficking assay

The pCD8a-eGFP plasmid (Addgene #86051) was digested with *EcoRI*/*NotI* to remove eGFP, and annealed oligos encoding the HA epitope were cloned to make pCD8a-HA. Chimeric CD8-based reporter plasmids were then constructed in pCD8a-HA, containing the CD8a ectodomain (residues 1-182) along with the transmembrane (TM) and C-tail of AAVR (residues 933-1049), AAVR2 (residues 1300-1380), FURIN (residues 716-794) or CI-MPR (residues 2305-2491), which were synthesised as Geneblocks (Integrated DNA Technologies, IDT, IA, USA) with *EcoRI*/*MluI* ends. Chimeric AAVR and AAVR2 C-tail mutants were similarly constructed by cloning Geneblocks (IDT) in pCD8-AAVR-HA and pCD8-AAVR2-HA respectively digested with *EcoRI*/*MluI* (see **Table S1** for sequence information). HEK293T cells (3.5×10^5^) were seeded in a 6-well plate overnight in normal growth medium. Cells were then transfected with CD8-based reporter plasmids using calcium phosphate precipitation. After 48 h, cells were collected, washed with FACS buffer (2% (v/v) FCS/PBS) and stained on ice with CD8-APC-Vio770-conjugated mouse antibody (1:100, Miltenyi, Germany; 130-110-681) for 1 h. Cells were then washed 3x with FACS buffer and then fixed and permeabilised with CytoFix (BD Biosciences, CA, USA; 554655) for 20 min on ice. Cells were then incubated with HA-PE-conjugated rabbit antibody (1:100; Abcam; ab72564) for 1 h on ice, then washed twice with CytoWash (BD Biosciences; 560105), before analysis by flow cytometry (BD Fortessa).

### Lentivirus-mediated overexpression of AAVR, AAVR2 and miniAAVR2

AAVR, AAVR2 and miniAAVR2 were overexpressed using lentiviral vectors. First, cDNAs encoding AAVR, AAVR2 (Horizon Discovery #BC0316172 & #BC045624) and miniAAVR2 (IDT) were cloned into a FUW lentiviral plasmid backbone (Addgene #14882). The plasmids were then co-transfected with pRSV-Rev (Addgene #12253), pMDLg/pRRE (Addgene #12251) and pMD2.G (Addgene #12259) using calcium phosphate into HEK293T cells. The supernatant containing virus was filtered and resuspended in FBS/media and snap frozen for storage. For lentiviral transduction, cells were spinoculated with lentiviruses in the presence of 8 μg/mL polybrene at 1,500 rpm for 90 min.

### Analysis of post-translational modifications (PTMs)

PTMs in AAVR and AAVR2 were analyzed from the data collated from PhosphositePlus. PTMs included phosphorylation (serine, threonine and tyrosine), lysine ubiquitylation, lysine acetylation, arginine methylation and O-linked glycosylation.

### Surface expression of AAVR and AAVR2

AAVR-or AAVR2-overexpressing HuH-7 cells were collected, washed three times with PBS containing 4% (w/v) BSA and incubated with anti-AAVR (1:200) or anti-AAVR2 (1:200) antibodies at RT for 1 h. After three further PBS washes, the following secondary antibodies were added: Alexa Fluor 488-conjugated Goat anti-Mouse (1:1,000; Invitrogen; A11001) for AAVR or Goat anti-Rabbit (1:1,000; Invitrogen; A11034) for AAVR2 and incubated at RT (1 h). Cells were then collected, washed and suspended in PBS containing DAPI (1:1,000; Invitrogen; D1306). Flow cytometry was performed to quantify the expression of DAPI (live/dead staining) and Alexa Fluor 488 signals. Data were analyzed using FlowJo software (Becton, Dickinson & Company, OR, USA).

### AAV transduction

AAV vectors expressing eGFP (Addgene #83279) were double cesium chloride-purified and titred by the Vector and Genome Engineering Facility (VGEF, Westmead, Australia). Unless otherwise stated, 50,000 cells were seeded in a 24-well plate followed by the addition of AAV after 24 h. Flow cytometry was performed 48-72 h later to quantify the percentage of GFP-positive cells and used as an indicator of transduction. Data analysis was performed with FlowJo software.

### Generation of AAVR and AAVR2 knockout cells

The generation of *AAVR* and *AAVR2* KO eHAP and HeLa cells using CRISPR/Cas9 has been described previously^10^. To generate HEK293T *AAVR* and *AAVR2* KO cells, single-guide RNAs (sgRNAs) targeting exon 2 of *AAVR* and exon 1 of *AAVR2* acquired as single-stranded oligonucleotides (Integrated DNA Technologies) and cloned into the pLKO.1-puro U6 sgRNA BfuA1 plasmid (modified with H2B-2A-mCherry from Addgene plasmid #52628). HEK293T cells were transiently transfected with sgRNA-expressing plasmids and pLV-UbC-eGFP2aCas9 (Addgene #53190). Single cells co-expressing eGFP and mCherry were collected with fluorescence-activated cell sorting (FACS) in 96-well plates and expanded. Knockout clones were validated with Sanger sequencing and westerns blots. Sequences of sgRNAs used are listed in **Table S1**.

### RNA extraction, cDNA synthesis and Quantitative real-time PCR

TRIzol reagent (Invitrogen; 15596026) was used to isolate total RNA following the manufacturer’s instructions and iScript cDNA synthesis kit (Bio-Rad, CA, USA; 1725034) was used to synthesize complementary DNA (cDNA). Expression of *AAVR* and *AAVR2* were measured using the SensiFast MasterMix (Bioline, London, UK; 84005) on a CFX96 PCR machine. The PCR condition used was as follows: 95 °C for 2 min, 35 cycles of 95 °C for 5 s, 60 °C for 10 s and 72 °C for 15 s. The β2-microglobulin (β*2M*) was used for normalization and the relative expression of *AAVR* and *AAVR2* was calculated using the ddCT method (primers are listed in **Table S1**).

### Immunofluorescence

For steady-state immunofluorescence analysis of AAVR and AAVR2, HuH-7 cells (4×10^3^) were seeded on Ibidi 12-well slides (ibidi, Germany; 81201). After 24 h, cells were fixed in 4% (w/v) formaldehyde (ThermoFisher; 28906), permeabilised with 0.2% (v/v) Triton X-100 in PBS and blocked with 3% (w/v) BSA in Triton X-100/PBS. Primary antibody staining was then performed with mouse anti-AAVR, rabbit anti-AAVR2 or rabbit anti-GM130 (Abcam; ab52649) antibodies overnight at 4 °C. Cells were centrifuged, washed with PBS and stained with appropriate secondary antibodies for 1 h: anti-rabbit IgG Alexa Fluor 488 (ThermoFisher; A11034), anti-mouse IgG Alexa Fluor 488 (ThermoFisher; A11001), anti-rabbit IgG Alexa Fluor 594 (ThermoFisher; A11020) or anti-mouse IgG Alexa Fluor 594 (ThermoFisher; A11012). Nuclear staining was done with DAPI. Slides were mounted with VECTASHIELD (Vector Laboratories, CA, USA; H100010) and imaged using a Leica SP8 confocal microscope (Leica Microsystems, Wetzlar, Germany).

For analysis, eHAP cells overexpressing AAVR or AAVR2 (5×10^4^) were seeded on chambered glass coverslips (Lab-Tek, ThermoFisher, 155411) overnight. AAV8 (100,000 vg/cell) was added to the cells and incubated for either 1 h at 4 °C (attachment) or 3 h at 37 °C (entry). Cells were fixed with 4% (w/v) formaldehyde for 15 min at RT, washed with PBS and incubated with mouse anti-AAV8 (Progen, Heidelberg, Germany; 692318) and rabbit anti-HA (Abcam; Ab9110) antibodies followed by three rounds of washing with PBS and incubation with appropriate secondary antibodies: anti-mouse IgG (Jackson Lab; 715-005-150) or anti-rabbit IgG (Jackson Lab; 711-005-152) conjugated to fluorophores Alexa647-NHS (ThermoFisher; A20006) or Cy3B-NHS (GE Healthcare; GEPA13004). For staining F-actin, an Alexa488 conjugated Phalloidin was used (ThermoFisher; A12379). Total International Reflection Fluorescence (TIRF) imaging was performed using an Olympus (now known as Evident) cellTIRF-4Line System equipped on an Olympus IX83 microscope frame with a 100X oil immersion TIRF objective (NA 1.49, Olympus, Japan) and an Evolve EMCCD camera (Photometrics, Arizona, US). Laser lines 488, 561 and 647 nm (100 mW, Coherent, US) were used for the excitation of fluorophores with a 405/488/561/647 nm Laser Quad Band Set (Chroma, CA, US). Images were acquired using CellSens software (Version 4.1.1, Olympus, Japan). Image post-processing including look-up-table and co-localization calculations were performed using ImageJ (National Institutes of Health, US) and Matlab (ver2021a, MathWorks, Inc. US) as previously described^29^.

### Generation of HeLa cells stably expressing miniAAVR2

MiniAAVR2 cDNA was cloned in a FUW lentiviral vector backbone (Addgene #14882) modified to express mCherry or mCherry and miniAAVR2 separated by the self-cleaving P2A peptide for co-expression. HeLa cells (5×10^5^) were transduced with lentiviruses expressing mCherry-P2A-miniAAVR2 (HeLa-miniAAVR2) or mCherry alone (HeLa-EV). Cells positive for mCherry were enriched with FACS and maintained in standard media for experiments or stored for future use.

### AAV Neutralization assay

AAV8-eGFP (40 μL, 5×10^8^ vg/μL) was mixed with the following concentrations of anti-AAV8 (ADK8/9, Progen, 651161): 500 ng, 250 ng, 125 ng and 62.5 ng (40 μL) and incubated for 1 h at 37 °C to facilitate neutralization. The mixture (80 μL) was added to HeLa-EV-or miniAAVR2-expressing cells (50,000) and incubated for 48 h before the quantification of transduction using flow cytometry.

## Statistical analysis

All experiments were conducted three times unless otherwise stated and data represented as mean ± SD. Welch’s t-test was used for statistical analysis using GraphPad Prism where ns=not significant, \**p*<0.05, \*\**p*<0.01, \*\*\**p*<0.001 and \*\*\*\**p*<0.0001.

## Data availability statement

All raw mass spectrometry data have been deposited to the ProteomeXchange Consortium via the PRIDE partner repository with the dataset identifier PXD074622. Requests for other data should be made to the corresponding author.

## Acknowledgements

We acknowledge funding support from the OHMR NSW Health Cell & Gene Therapy Grant (JEJR & CGB), National Health and Medical Research Council (NHMRC) Ideas Grants #2037697 and #2047345 (both to CGB & BPD), NHMRC Investigator grant #1177305 (JEJR), Brandon Capital CUREator (JEJR, BPD & CGB), Therapeutic Innovation Australia (BPD & CGB) and Cure the Future Foundation for philanthropic support. The authors thank Sydney Cytometry, Sydney Mass Spectrometry at University of Sydney and Centenary Institute for their support.

## Author contributions

Conceptualization, BPD, JEJR and CGB; data curation, BPD, MST and CGB; formal analysis, BPD, QPS and CGB; methodology, BPD, RN, JV, YF, CM, VK, CC, DG, AS, QPS, MST and CGB; supervision, BPD and CGB; visualization, BPD, MST, QPS and CGB; writing - original draft, BPD, QPS and CGB; writing - review and editing, BPD, JEJR and CGB; funding acquisition, BPD, JEJR and CGB.

## Supplementary Information

**Fig. S1 (related to Fig. 1):**
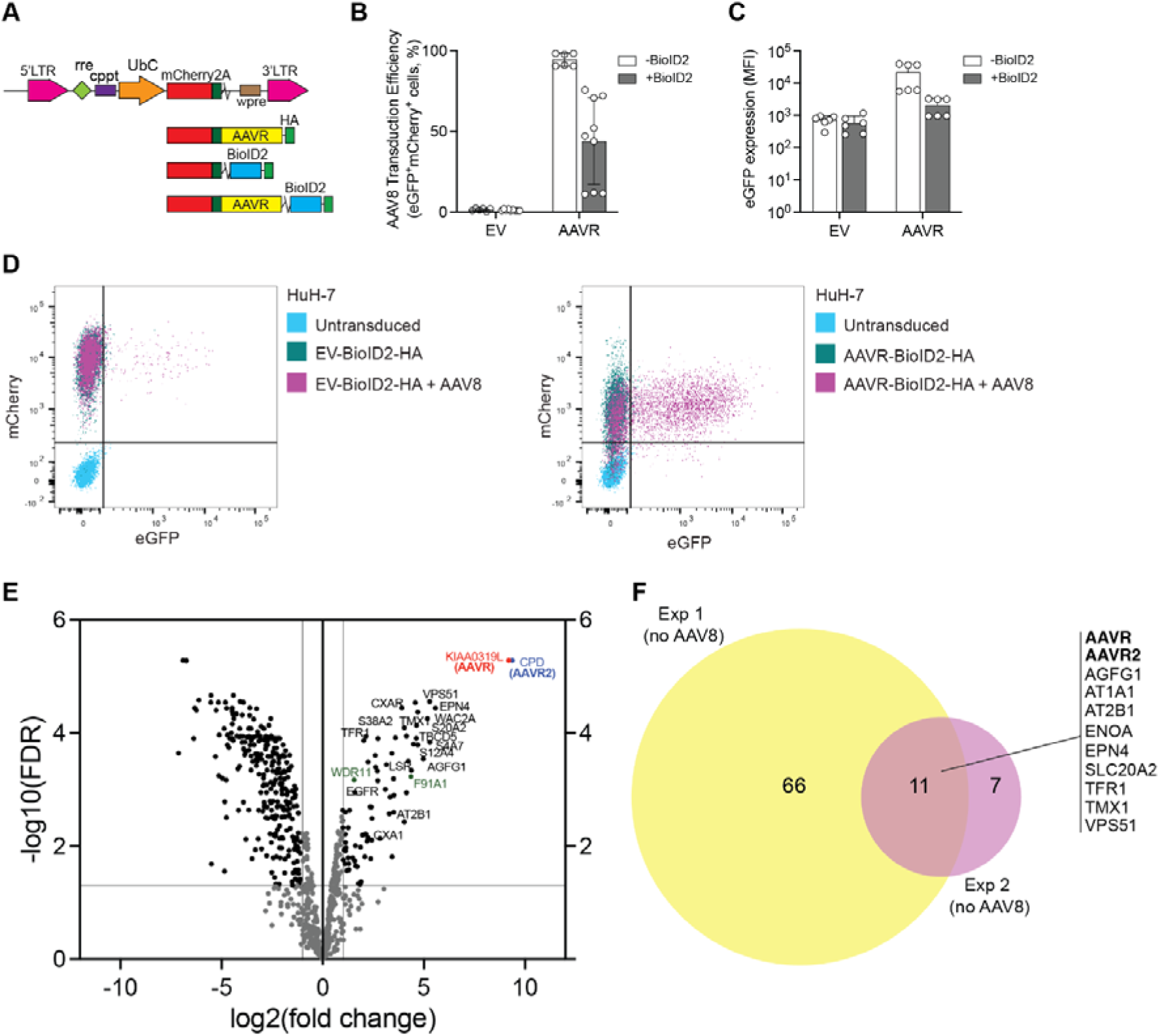
Reagent validation and AAVR BioID performed in the absence of AAV8. (**A**) Schematic of lentiviral vectors used to determine whether the BioID2 tag on AAVR C-term impacts AAV8 transduction: mCherry2A alone (empty vector, EV), mCherry-2A-BioID2 (EV-BioID2), mCherry-2A-AAVR (AAVR) and mCherry-2A-AAVR-BioID2 (AAVR-BioID2). (**B-C**) HEK293T *AAVR* KO cells were transduced with lentiviral vectors for 48 h, then transduced with AAV8-eGFP (100,000 vg/cell) for an additional 48 h. Flow cytometric analysis was used to measure eGFP expression in mCherry-positive cells. AAV8 transduction (**B**) and eGFP expression measured by mean fluorescence intensity (MFI) (**C**) are shown. Data in **B-C** represent the mean ± SD of 2-3 independent experiments performed in triplicate. (**D**) Representative flow cytometric overlay plots of HuH-7 cells subjected to biotin labelling (16 h) and AAV8 transduction (5,000 vg/cell, 30 min) followed by analysis after 48 h after removal of AAV8. EV-BioID2-HA (*left panel*) and AAVR-BioID2-HA (*right panel*) are shown. The x-& y-axes represent eGFP and mCherry expression respectively. (**E**) Volcano plot of enriched proteins after AAVR BioID performed in HuH-7 cells in the absence of AAV8. (**F**) Venn diagram summarising two independent AAVR BioID experiments (Exp 1 and Exp 2); proteins enriched in both experiments are listed (see also **Supplementary data**).

**Fig. S2 (related to Fig. 2):**
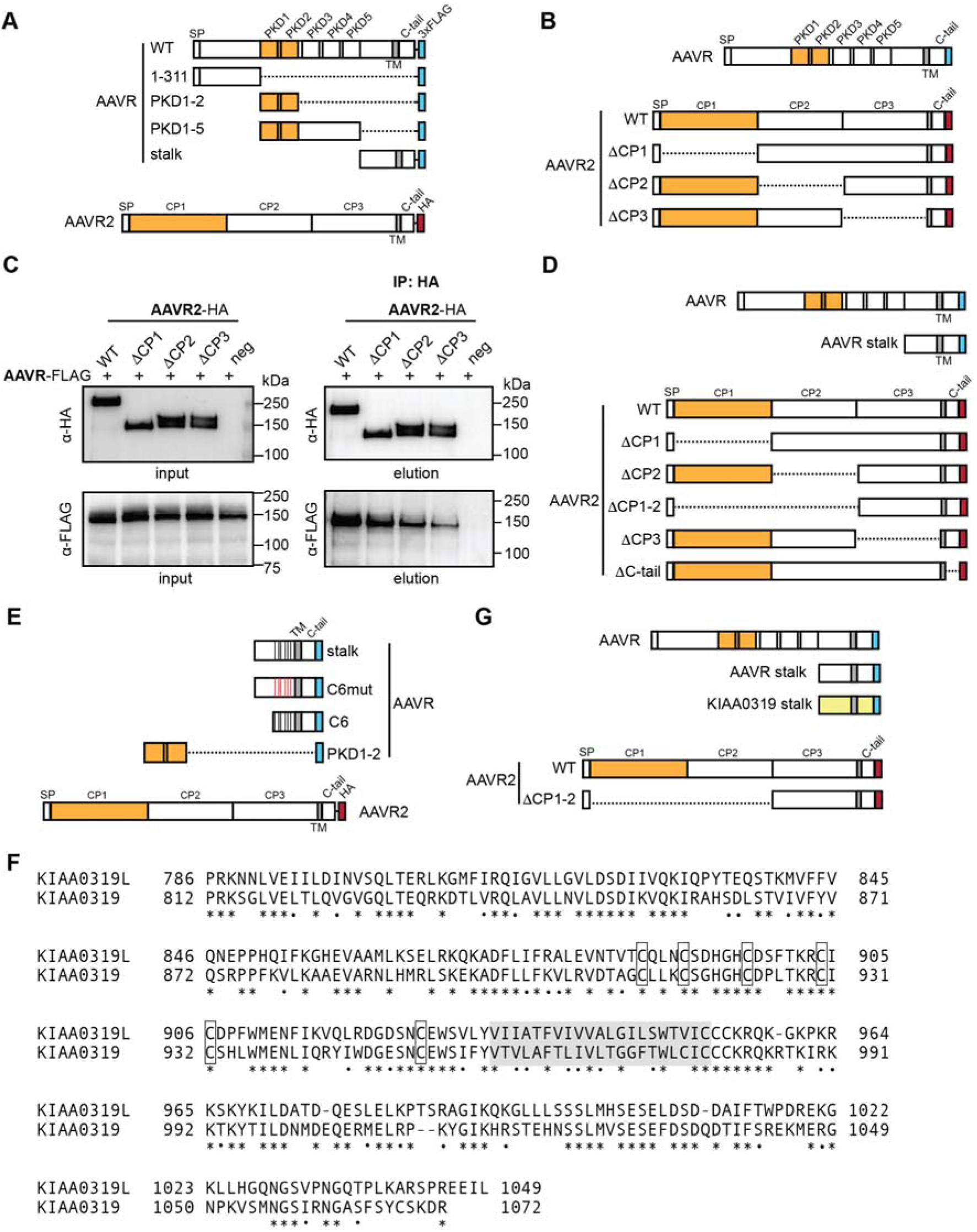
Refinement of the minimal interacting regions of AAVR and AAVR2. (**A**) Schematic of AAVR-3xFLAG deletion mutants tested for interaction with AAVR2-HA using co-immunoprecipitation (Co-IP). (**B**) Schematic of AAVR2-HA deletion mutants tested for a direct interaction with AAVR-3xFLAG. (**C**) Mapping the minimal region in AAVR2 interacting with AAVR by Co-IP using HA beads for pulldown. Input and eluate blots were loaded separately and each probed with anti-HA and anti-FLAG antibodies; Δ represents deletion and neg represents untransfected cells used as negative control. Protein markers on all blots indicate molecular weight in kDa. (**D**) Schematic showing AAVR2 deletion mutations used in Co-IP experiments with the AAVR stalk region. (**E**) Schematic of the AAVR stalk region and mutants tested for interaction with WT AAVR2. (**F**) Protein alignment of AAVR (KIAA0319L) and KIAA0319 stalk regions showing amino acid identity. Cysteine residues in the EGF-like domain are boxed; TM region is shaded in grey. In **A**, **B**, **D**, **E** & **G**, the AAV-interacting regions in AAVR and AAVR2 are highlighted in orange. (**G**) Schematic of the AAVR stalk region and stalk region in KIAA0319 tested for binding with WT AAVR2 and CP1 and CP2 deleted mutant (ΔCP1-2).

**Fig. S3 (related to Fig. 3):**
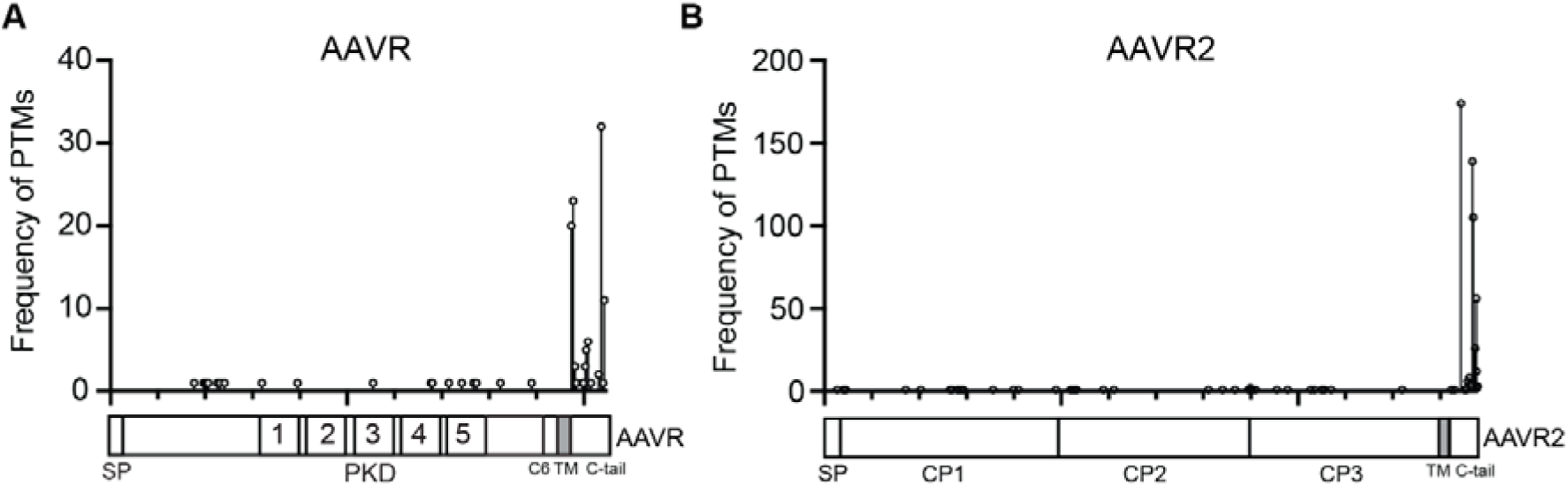
Post-translational modifications (PTMs) in AAVR and AAVR2. Data was collated from PhosphositePlus and depicted on the schematics for AAVR (**A**) and AAVR2 (**B**). The frequency of PTMs is depicted on the y-axis; schematics are drawn to scale.

**Fig. S4 (related to Fig. 4):**
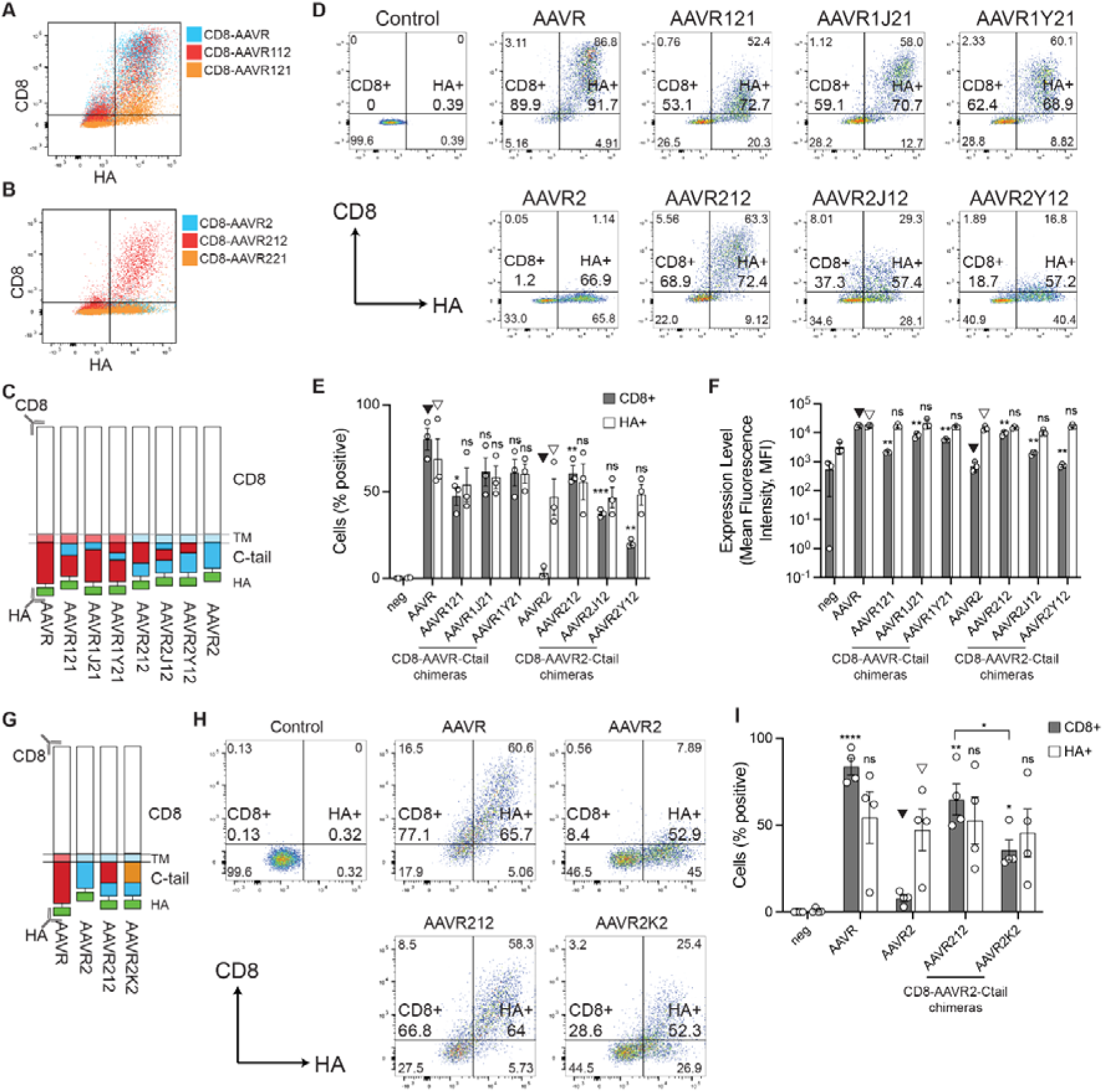
**Refinement of AAVR and AAVR2 C-tail motifs that govern cell surface expression. (A-B**) Overlays of representative flow cytometry plots showing CD8-AAVR,-AAVR112 and-AAVR121 (**A**), CD8-AAVR2,-AAVR212 and-AAVR221 (**B**); x-axis=HA, y-axis=CD8 (the CD8+ gate includes quadrants Q1 and Q2; the HA+ gate includes quadrants Q2 and Q3). (**C**) Schematic of CD8 fusion reporter constructs comparing different chimeric AAVR and AAVR2 C-tails resulting from domain swapping. Full details of amino acid co-ordinates are in **Table S2**. Unmodified AAVR and AAVR2 TMs and C-tails were used as controls. (**D**) Representative flow cytometry plots (y-axis=CD8, x-axis=HA). (**E-F**) Summary of flow cytometric analysis of CD8 and HA expression depicting cells with CD8 surface expression (%) and total HA expression (%) (**E**), and expression level (MFI) of CD8 and HA (**F**). Arrowheads indicate the sample used for each statistical comparison for CD8+ and HA+ samples (*n*=3). (**G**) Schematic of CD8 fusion reporter constructs comparing different chimeric C-tails from AAVR, AAVR2 and KIAA0319. (**H**) Representative flow cytometry plots (y-axis=CD8, x-axis=HA). (**I**) Summary of flow cytometric analysis of CD8 and HA expression depicting cells with CD8 surface expression (%) and total HA expression (%) (*n*=4). Arrowheads indicate the sample used for each statistical comparison for CD8+ and HA+ samples. Welch’s t-test was used for statistical analysis where ns=not significant, \**p*<0.05, \*\**p*<0.01, \*\*\**p*<0.001 and \*\*\*\**p*<0.0001.

**Fig. S5 (related to Fig. 5):**
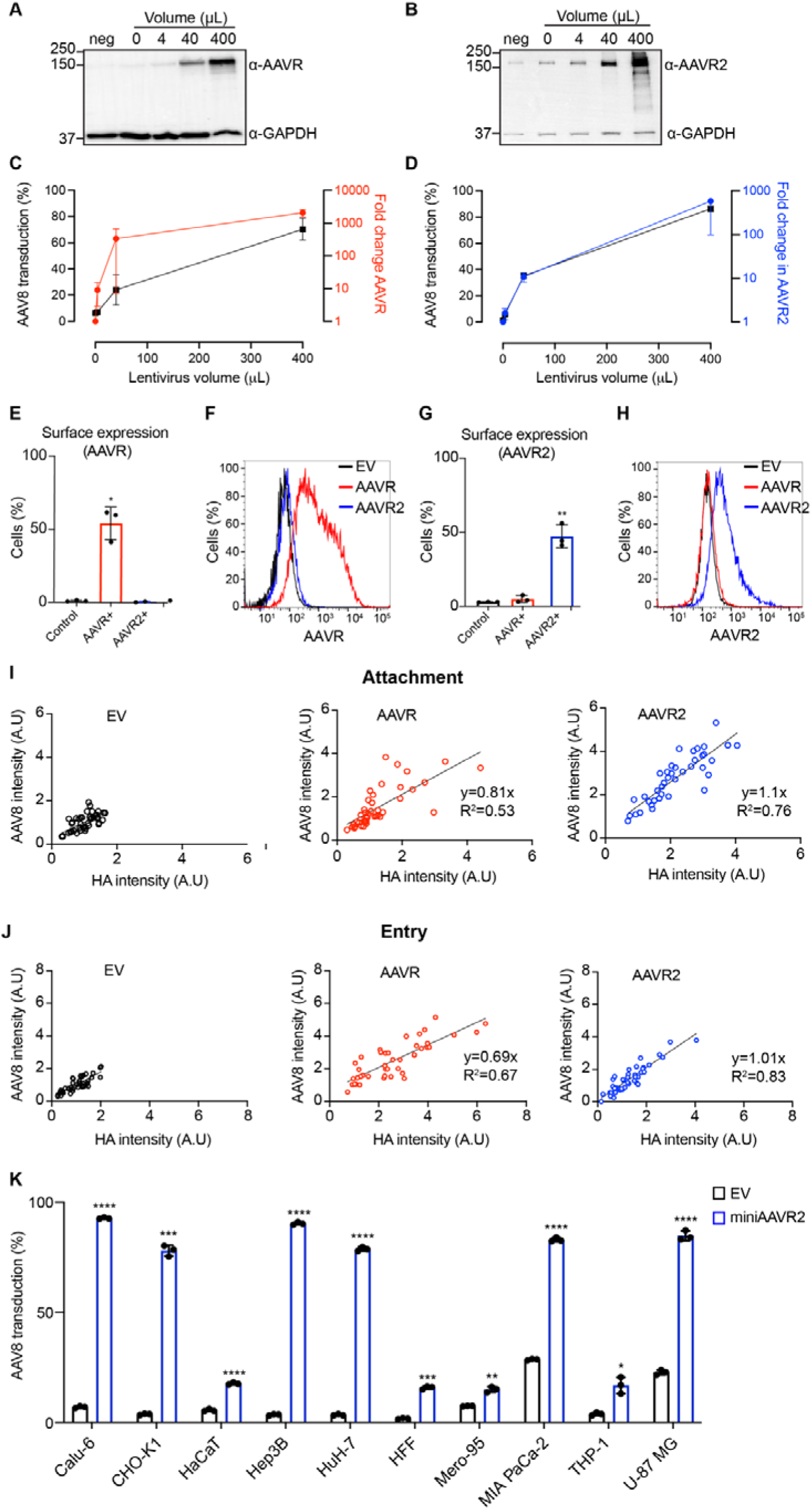
AAVR or AAVR2 overexpression increases AAV transduction in a serotype-specific manner. (**A**) Representative western blots performed with anti-AAVR or anti-GAPDH antibodies in HuH-7 cells transduced with indicated volumes of AAVR-expressing lentiviral vectors. neg=untransduced cells. (**B**) Representative western blots performed with anti-AAVR2 or anti-GAPDH antibodies in HuH-7 cells transduced with indicated volumes of lentiviral vectors to overexpress AAVR2. (**C**) Plot showing the relationship between AAV8 transduction and AAVR expression in HuH-7 cells (*n*=2). (**D**) Plot showing the relationship between AAV8 transduction and AAVR2 expression in HuH-7 cells (*n*=3). (**C-D**) X-axis: volume of lentiviral vector expressing AAVR or AAVR2, left y-axis: AAV8 transduction measured with flow cytometry and right y-axis: expression of AAVR or AAVR2 measured with quantitative real-time PCR. (**E**) Percentage of HuH-7 cells expressing AAVR at the surface following transduction with lentiviral vectors (400 µL) expressing AAVR or AAVR2 measured with flow cytometry. Cells transduced with empty vector (EV) were used as control (*n*=3). (**F**) Representative flow cytometry overlay plots showing the surface expression of AAVR in control, AAVR-overexpressing or AAVR2-overexpressing HuH-7 cell lines. The intensity of AAVR expression quantified using antibody-based flow cytometry (x-axis); y-axis represents the percentage of positive cells normalized to mode. (**G**) Cell surface expression of AAVR2 in EV, AAVR-or AAVR2-overexpressing HuH-7 cells. Overexpression was achieved with lentiviral vectors and EV transduced cells were used as control (*n*=3). (**H**) Representative flow cytometry overlay plots showing the surface expression of AAVR2 in control, AAVR-or AAVR2-overexpressing HuH-7 cell lines. (**I-J**) Plots showing the intensity of immunofluorescence staining for HA (representing AAVR or AAVR2) in x-axis and AAV8 in y-axis in control, AAVR-overexpressing and AAVR2-overexpressing eHAP cells at AAV8 attachment (**I**) and entry (**J**) conditions; A.U = arbitrary units. Overexpression was achieved using lentiviral vectors and EV transduced cells were used as controls. Each dot represents a single cell (*n*=42), and a simple linear regression curve was fitted using GraphPad Prism. (**K**) Transduction of AAV8 in a panel of cell lines overexpressing miniAAVR2 measured with flow cytometry. Overexpression was achieved using lentiviruses and EV transduced cells were used as control (*n*=3). (**E, G & K**) Welch’s t-test was used to determine the statistical significance. \**p*<0.05, \*\**p*<0.01, \*\*\**p*<0.001, \*\*\*\**p*<0.0001.

---

**Table S1:**
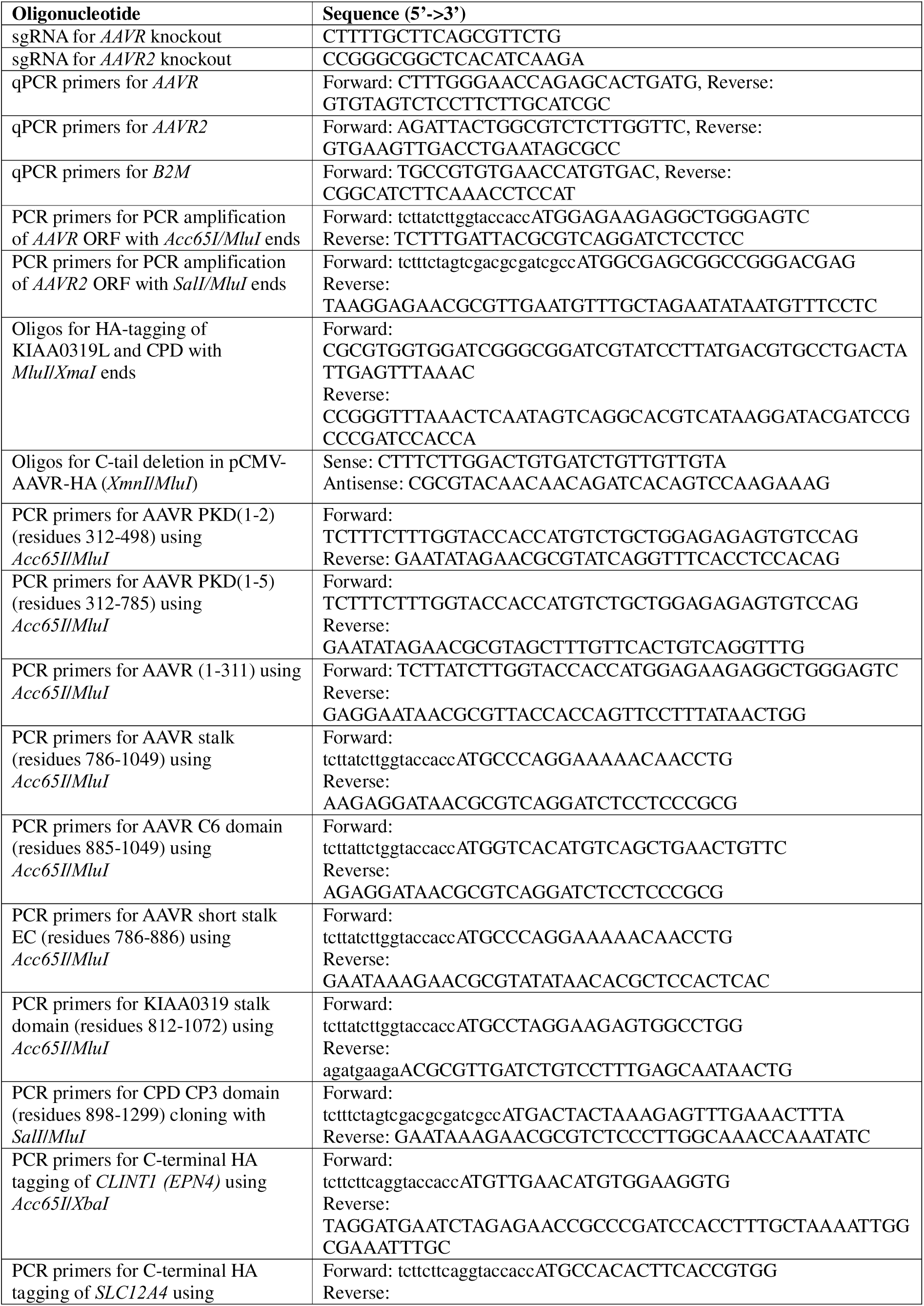
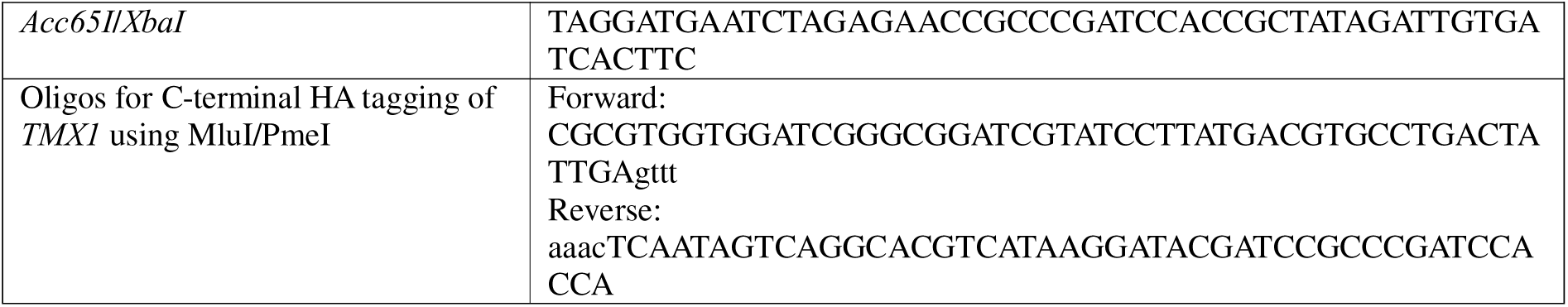
List of all oligonucleotides used in this study.

**Table S2:**
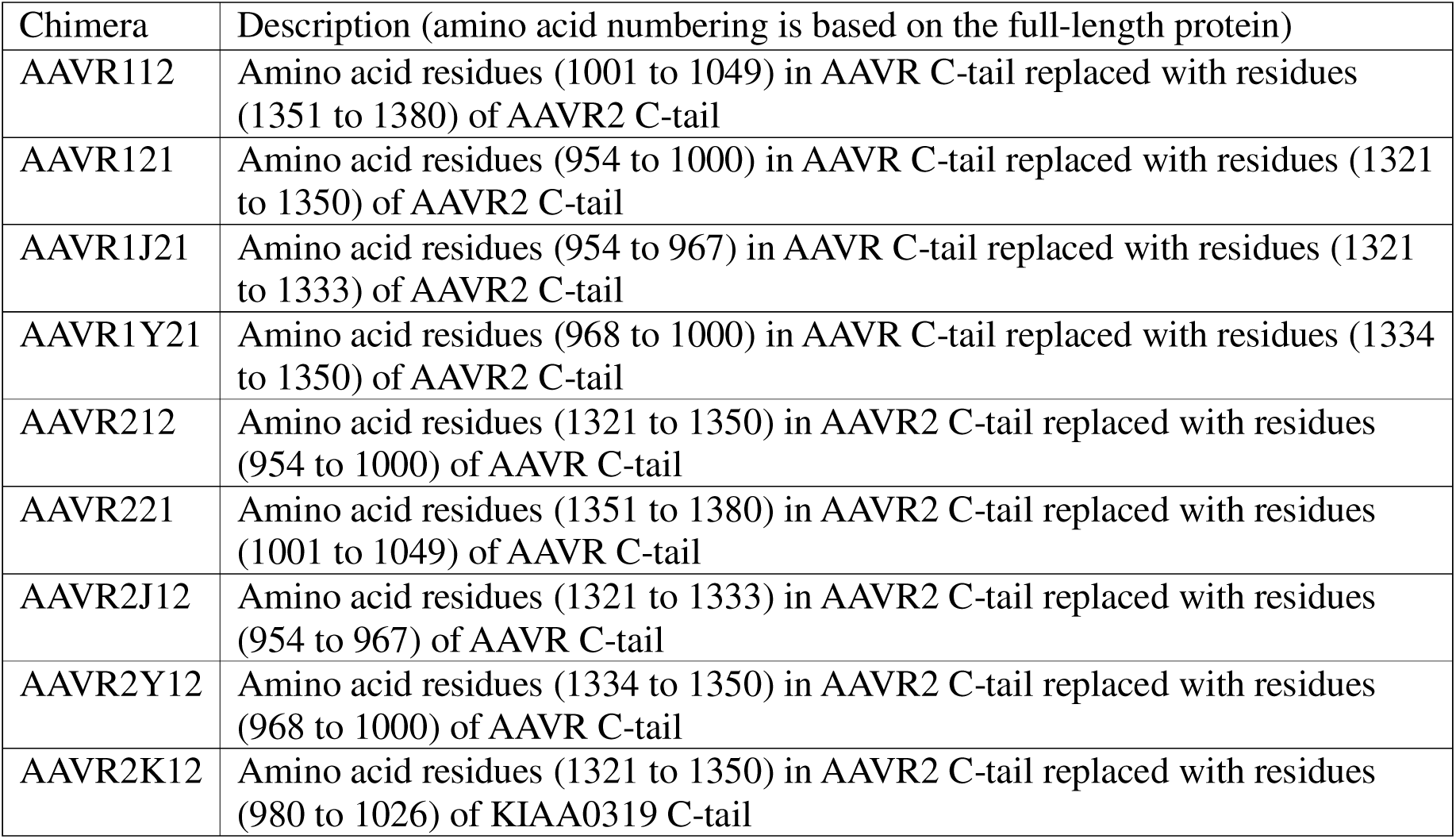
Details of chimeric AAVR/AAVR2 C-tail mutants used in CD8 reporter plasmids in this study.

## References

1. Wang, J.H., Gessler, D.J., Zhan, W., Gallagher, T.L., and Gao, G. (2024). Adeno-associated virus as a delivery vector for gene therapy of human diseases. Signal Transduct Target Ther 9, 78. 10.1038/s41392-024-01780-w.

2. Dhungel, B.P., Winburn, I., Pereira, C.D.F., Huang, K., Chhabra, A., and Rasko, J.E.J. (2024). Understanding AAV vector immunogenicity: from particle to patient. Theranostics 14, 1260–1288. 10.7150/thno.89380.

3. Duan, D. (2023). Lethal immunotoxicity in high-dose systemic AAV therapy. Mol Ther 31, 3123–3126. 10.1016/j.ymthe.2023.10.015.

4. Lek, A., Wong, B., Keeler, A., Blackwood, M., Ma, K., Huang, S., Sylvia, K., Batista, A.R., Artinian, R., Kokoski, D., et al. (2023). Death after High-Dose rAAV9 Gene Therapy in a Patient with Duchenne’s Muscular Dystrophy. N Engl J Med 389, 1203–1210. 10.1056/NEJMoa2307798.

5. Shen, W., Liu, S., and Ou, L. (2022). rAAV immunogenicity, toxicity, and durability in 255 clinical trials: A meta-analysis. Front Immunol 13, 1001263. 10.3389/fimmu.2022.1001263.

6. Nisanov, A.M., Rivera de Jesús, J.A., and Schaffer, D.V. (2025). Advances in AAV capsid engineering: Integrating rational design, directed evolution and machine learning. Mol Ther 33, 1937–1945. 10.1016/j.ymthe.2025.03.056.

7. Dhungel, B.P., Bailey, C.G., and Rasko, J.E.J. (2021). Journey to the Center of the Cell: Tracing the Path of AAV Transduction. Trends Mol Med 27, 172–184. 10.1016/j.molmed.2020.09.010.

8. Meyer, N.L., and Chapman, M.S. (2022). Adeno-associated virus (AAV) cell entry: structural insights. Trends Microbiol 30, 432–451. 10.1016/j.tim.2021.09.005.

9. Pillay, S., Meyer, N.L., Puschnik, A.S., Davulcu, O., Diep, J., Ishikawa, Y., Jae, L.T., Wosen, J.E., Nagamine, C.M., Chapman, M.S., and Carette, J.E. (2016). An essential receptor for adeno-associated virus infection. Nature 530, 108–112. 10.1038/nature16465.

10. Dhungel, B.P., Xu, H., Nagarajah, R., Vitale, J., Wong, A.C.H., Gokal, D., Feng, Y., Tabar, M.S., Metierre, C., Parsania, C., et al. (2025). An alternate receptor for adeno-associated viruses. Cell 188, 4924–4935.e4923. 10.1016/j.cell.2025.06.026.

11. Havlik, L.P., Das, A., Mietzsch, M., Oh, D.K., Ark, J., McKenna, R., Agbandje-McKenna, M., and Asokan, A. (2021). Receptor Switching in Newly Evolved Adeno-associated Viruses. J Virol 95, e0058721. 10.1128/JVI.00587-21.

12. Dudek, A.M., Pillay, S., Puschnik, A.S., Nagamine, C.M., Cheng, F., Qiu, J., Carette, J.E., and Vandenberghe, L.H. (2018). An Alternate Route for Adeno-associated Virus (AAV) Entry Independent of AAV Receptor. J Virol 92, e02213–02217. 10.1128/JVI.02213-17.

13. Pillay, S., Zou, W., Cheng, F., Puschnik, A.S., Meyer, N.L., Ganaie, S.S., Deng, X., Wosen, J.E., Davulcu, O., Yan, Z., et al. (2017). AAV serotypes have distinctive interactions with domains of the cellular receptor AAVR. J Virol 91, e00391–00317. 10.1128/JVI.00391-17.

14. Kim, D.I., Jensen, S.C., Noble, K.A., Kc, B., Roux, K.H., Motamedchaboki, K., and Roux, K.J. (2016). An improved smaller biotin ligase for BioID proximity labeling. Mol Biol Cell 27, 1188–1196. 10.1091/mbc.E15-12-0844.

15. Samavarchi-Tehrani, P., Samson, R., and Gingras, A.C. (2020). Proximity Dependent Biotinylation: Key Enzymes and Adaptation to Proteomics Approaches. Mol Cell Proteomics 19, 757–773. 10.1074/mcp.R120.001941.

16. Deng, H., Jia, G., Li, P., Tang, Y., Zhao, L., Yang, Q., Zhao, J., Wang, J., Tu, Y., Yong, X., et al. (2024). The WDR11 complex is a receptor for acidic-cluster-containing cargo proteins. Cell 187, 4272–4288.e4220. 10.1016/j.cell.2024.06.024.

17. Marino, S.M., and Gladyshev, V.N. (2012). Analysis and functional prediction of reactive cysteine residues. J Biol Chem 287, 4419–4425. 10.1074/jbc.R111.275578.

18. Pandey, K.N. (2010). Small peptide recognition sequence for intracellular sorting. Curr Opin Biotechnol 21, 611–620. 10.1016/j.copbio.2010.08.007.

19. Bonifacino, J.S., and Traub, L.M. (2003). Signals for sorting of transmembrane proteins to endosomes and lysosomes. Annu Rev Biochem 72, 395–447. 10.1146/annurev.biochem.72.121801.161800.

20. Cattin-Ortolá, J., Kaufman, J.G.G., Gillingham, A.K., Wagstaff, J.L., Peak-Chew, S.Y., Stevens, T.J., Boulanger, J., Owen, D.J., and Munro, S. (2024). Cargo selective vesicle tethering: The structural basis for binding of specific cargo proteins by the Golgi tether component TBC1D23. Sci Adv 10, eadl0608. 10.1126/sciadv.adl0608.

21. Pei, X., He, T., Hall, N.E., Gerber, D., Samulski, R.J., and Li, C. (2018). AAV8 virions hijack serum proteins to increase hepatocyte binding for transduction enhancement. Virology 518, 95–102. 10.1016/j.virol.2018.02.007.

22. Wallen, A.J., Barker, G.A., Fein, D.E., Jing, H., and Diamond, S.L. (2011). Enhancers of adeno-associated virus AAV2 transduction via high throughput siRNA screening. Mol Ther 19, 1152–1160. 10.1038/mt.2011.4.

23. Zhang, X., Hao, S., Feng, Z., Ning, K., Kuz, C.A., McFarlin, S., Richart, D., Cheng, F., Zhang-Chen, A., McFarlane, R., et al. (2024). Identification of SLC35A1 as an essential host factor for the transduction of multi-serotype recombinant adeno-associated virus (AAV) vectors. bioRxiv10.1101/2024.10.16.618764.

24. Dudek, A.M., Zabaleta, N., Zinn, E., Pillay, S., Zengel, J., Porter, C., Franceschini, J.S., Estelien, R., Carette, J.E., and Zhou, G.L. (2019). GPR108 is a highly conserved AAV entry factor. Molecular Therapy 28, 367–381.

25. Tabar, M.S., Parsania, C., Giardina, C., Feng, Y., Wong, A.C.H., Metierre, C., Nagarajah, R., Dhungel, B.P., Rasko, J.E.J., and Bailey, C.G. (2025). Intrinsically Disordered Regions Define Unique Protein Interaction Networks in CHD Family Remodelers. FASEB J 39, e70632. 10.1096/fj.202402808RR.

26. Sharifi Tabar, M., Giardina, C., Feng, Y., Francis, H., Moghaddas Sani, H., Low, J.K.K., Mackay, J.P., Bailey, C.G., and Rasko, J.E.J. (2022). Unique protein interaction networks define the chromatin remodelling module of the NuRD complex. FEBS J 289, 199–214. 10.1111/febs.16112.

27. Cox, J., and Mann, M. (2008). MaxQuant enables high peptide identification rates, individualized p.p.b.-range mass accuracies and proteome-wide protein quantification. Nat Biotechnol 26, 1367–1372. 10.1038/nbt.1511.

28. Shah, A.D., Goode, R.J.A., Huang, C., Powell, D.R., and Schittenhelm, R.B. (2020). LFQ-Analyst: An Easy-To-Use Interactive Web Platform To Analyze and Visualize Label-Free Proteomics Data Preprocessed with MaxQuant. J Proteome Res 19, 204–211. 10.1021/acs.jproteome.9b00496.

29. Su, Q.P., Zhao, Z.W., Meng, L., Ding, M., Zhang, W., Li, Y., Liu, M., Li, R., Gao, Y.Q., Xie, X.S., and Sun, Y. (2020). Superresolution imaging reveals spatiotemporal propagation of human replication foci mediated by CTCF-organized chromatin structures. Proc Natl Acad Sci U S A 117, 15036–15046. 10.1073/pnas.2001521117.

